# Box C/D Small Nucleolar Ribonucleoproteins Regulate Mitochondrial Surveillance and Innate Immunity

**DOI:** 10.1101/2021.04.28.441759

**Authors:** Elissa Tjahjono, Alexey V. Revtovich, Natalia V. Kirienko

**Affiliations:** Department of BioSciences, 6100 Main St, MS140, Rice University, Houston TX 77005 USA

**Keywords:** mitochondria, snoRNPs, surveillance, translation inhibition, infection response

## Abstract

Monitoring of mitochondrial functions is crucial for organismal survival. This task is performed by mitochondrial surveillance or quality control pathways, which are activated by signals originating from mitochondria and relayed to the nucleus (retrograde response) to start the transcription of protective genes. In *Caenorhabditis elegans*, several systems exist, including the UPR^mt^, MAPK^mt^, and the ESRE pathway. These pathways are highly conserved and their loss results in compromised survival following mitochondrial stress.

In this study, we found a novel interaction between the box C/D snoRNA core proteins (snoRNPs) and mitochondrial surveillance and innate immunity pathways. We showed that C/D snoRNPs are required for the full expressions of UPR^mt^ and ESRE upon stress. Meanwhile, we found that the loss of C/D snoRNPs increased immune responses. Understanding the “molecular switch” mechanisms of interplay between these pathways may be important for understanding of multifactorial processes, including response to infection or aging.

## Introduction

All living organisms require the maintenance of cellular homeostasis very different than their surroundings. Maintaining these conditions requires constant surveillance for disruption and metabolic adjustments to reacquire the proper biochemical balance. Meanwhile, a variety of insults can disrupt this balance, ranging from environmental changes to pathogen infection to metabolic dysfunction. Indeed, almost all cellular pathways are disrupted in one infection or another, including protein translation (1, 2), the proteostatic machinery (3–5), the cytoskeleton (6), the endoplasmic reticulum (7, 8) and others (9, 10).

Given the central role of the mitochondria in energy production, biosynthesis of heme groups, lipid metabolism, the regulation of iron and calcium homeostasis, and production of reactive oxygen species (ROS), it should be no surprise that mitochondria are also impacted by disease and infection (11–13). Therefore, mitochondria are subjected to several important surveillance pathways. The two best known are the PINK1/Parkin axis for macroautophagic mitochondrial recycling (commonly known as mitophagy) and the unfolded protein response in mitochondria (UPR^mt^) (14–17). Both systems monitor the functionality of mitochondrial protein import. Failure to import PINK1 activates its kinase function, resulting in subsequent recruitment of autophagic machinery. Under the same compromised mitochondrial import conditions, rerouting of the key transcription factor ATFS-1/ATF5 to the nucleus activates the expression of chaperones and other stress mediators.

A third, rather more elusive, mitochondrial pathway that has been published utilizes the DLK-1/SEK-3/PMK-3 MAP kinase cascade (which we will refer to as the MAPK^mt^ pathway) was identified by activating mitochondrial stress and searching for differentially expressed genes that were independent of ATFS-1/ATF5 regulation (18). Regulation of this pathway appears to involve a C/EBP family transcription factor and senses disruption of the mitochondrial electron transport chain (ETC). It is involved in the extended lifespan observed in long-lived mitochondrial (Mit) mutants (18). Interestingly, fluvastatin, which disrupts mevalonate metabolism and prevents geranylgeranylation of certain components of the vesicular trafficking system, also activates the MAPK^mt^ pathway, indicating that surveillance of mitochondrial cholesterol metabolism is also important (19).

Our lab has previously identified a key mitochondrial surveillance program in *C. elegans* regulated by cellular ROS (20, 21). This pathway, known as the Ethanol and Stress Response (ESRE) network, is named after an 11-nucleotide motif found in the promoter region of hundreds of genes in *C. elegans* and ethanol-responsive genes in mice, and is activated by a range of abiotic triggers (22–24). Interestingly, exposure to the opportunistic human pathogen, *Pseudomonas aeruginosa*, which produces a xenobiotic siderophore called pyoverdine that hijacks mitochondria-resident iron from *C. elegans*, also activates the ESRE network (20, 25, 26). Active study from our lab and others has linked several determinants to ESRE gene expression, including the JumonjiC-domain containing protein JMJC-1/Riox1 (also known as NO66) (22), the PBAF nucleosome remodeling complex (27), and a family of bZIP transcription factors (ZIP-2, ZIP-4, CEBP-1, and CEBP-2) (20), and a Zn-finger transcription factor SLR-2 (22, 28). Importantly, the ESRE motif, the genes regulated by it, and their activation in response to stress are ancient and evolutionarily conserved from *C. elegans* to humans (22, 24) and appears to be the first known pathway to respond to intracellular reductive stress (21).

Mitochondrial surveillance programs not only activate programs to reacquire homeostasis, they also activate innate immune functions, a process sometimes termed surveillance immunity (29). However, innate immune activation is energetically costly and requires considerable energy conversion (30, 31) and excess immune activity is associated with a broad range of deleterious health outcomes. Thus, it behooves the organism to carefully balance the need to reacquire homeostasis and repair damage with stimulating innate immune functions that can cause further damage. How organisms navigate this choice remains a poorly understood area of biology.

Small nucleolar ribonucleoproteins (snoRNPs) are small complexes that catalyze modifications to RNA in cells. Generally speaking, there are two major groups of snoRNPs, called the box C/D and box H/ACA families, categorized based on their functions and the secondary structures of their snoRNA components (32). The core members of box C/D snoRNPs, consisting of FIB-1/Fibrillarin (the catalytic methyltransferase), NOL-56/Nop56, NOL-58/Nop58, and M28.5/SNU13 (**Figure 1A**), assist in site-specific 2’-*O*-methylation while box H/ACA family, consisting of Y66H1A.4/Gar1, Y48A6B.3/Nhp2, NOLA-3/Nop10, and K01G5.5/dyskerin, is involved in pseudouridylation (**Figure 1B**) (33). Both families target mostly ribosomal RNA, with the modifications typically clustered at biologically important locations (34). The snoRNA has sequence complementarity to the modification target site and serves as an aid to localize the enzymatic function of the snoRNP complex (34). snoRNP complexes had been suggested to have functional roles well beyond the processing of rRNA, including 2′-O-methylation, splicing, and translation of mRNAs (35–39).

**Figure 1.**
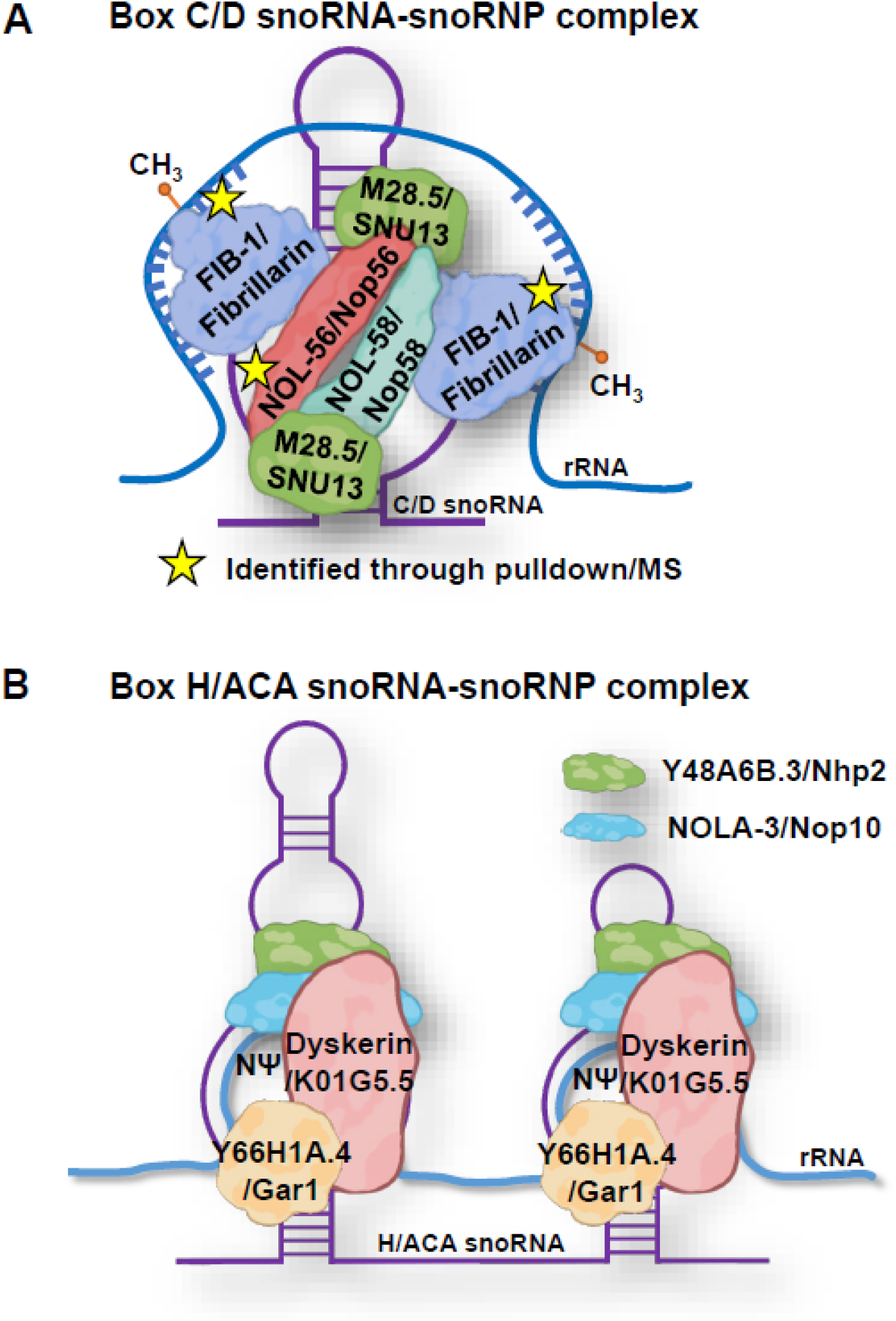
Cartoon representation of snoRNA and snoRNP complexes. **(A)** Box C/D snoRNA and C/D snoRNP core protein members, stars indicated proteins identified through oligo pulldown-mass spectrometry experiment. **(B)** Box H/ACA snoRNA and H/ACA snoRNP core protein members.

In this study, we identified a non-canonical role for box C/D snoRNPs where they appear to serve as a molecular switch that activates mitochondrial surveillance and represses conventional innate immune processes. For example, box C/D snoRNPs upregulate ESRE and UPR^mt^ while downregulating the function of the PMK-1/p38 MAPK pathway. Contrarily, knockdown of box C/D snoRNPs upregulated MAPK^mt^ pathway effectors, but this was likely a secondary effect from the loss of MAPK^mt^ repression by the UPR^mt^, which is characteristic of the complicated interactions between these surveillance systems. Since box C/D snoRNPs and these mitochondrial surveillance systems are all conserved between *C. elegans* and humans, our results may lead to a better understanding of processes affecting mitochondrial health and innate immune pathways in human diseases.

## Results

### Identification of FIB-1/Fibrillarin and NOL-56/Nop56 as regulators of the ESRE pathway

To identify additional regulatory components of the ESRE pathway, we used a biochemical pulldown method (**Figure 2**, **Figure 2-source data 1-4**). A 5’ biotinylated oligonucleotide comprised of a 3× or 4× tandem repeat of the consensus 11-nucleotide motif was used as bait. Young adult *C. elegans* were exposed to either DMSO, 1 mM phenanthroline (a chemical iron chelator), or 50 μM rotenone (inhibitor of electron transport chain Complex I) to trigger mitochondrial damage and ESRE gene activation (20). Proteins were extracted from the cytoplasm and nuclei and mixed with the biotinylated ESRE bait and then pulled down using streptavidin-coated magnetic beads. Electrophoretic mobility shift assay (EMSA) (27) was used to optimize enrichment of ESRE-binding proteins and to verify specificity (**Figure 2B**).

**Figure 2.**
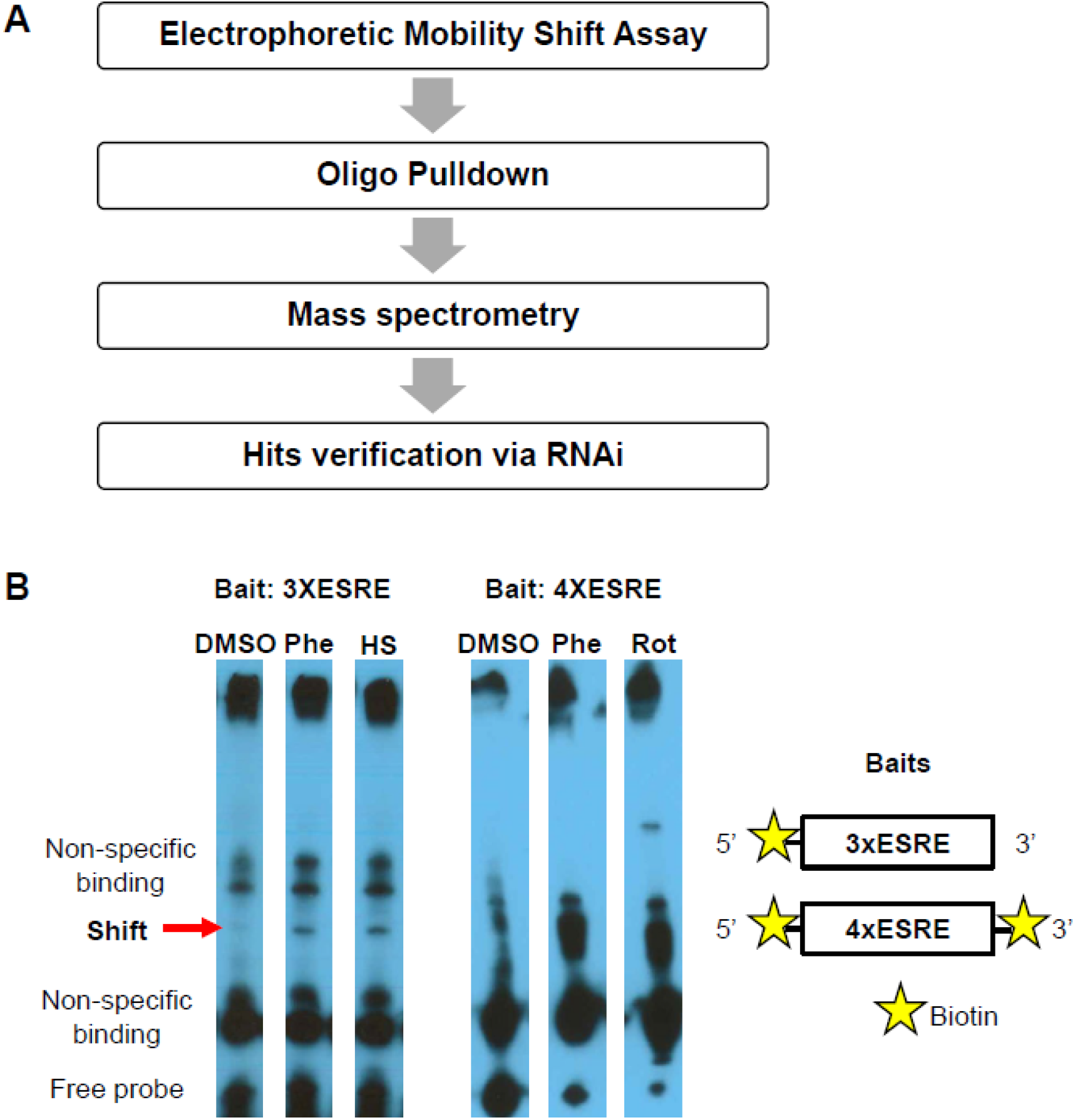
Proteomic assays revealed the presence of ESRE-binding factor(s). **(A)** Schematic of planned biochemical assays to identify ESRE regulators. **(B)** Electrophoretic mobility shift assay (EMSA) showed the presence of the ESRE-binding motif through the identified ‘Shift’.

We detected multiple bands when EMSA was performed with three tandem ESRE sequences (3XESRE) as a bait (**Figure 2B**, **Figure 2-source data 1-2**). These bands might indicate constitutive binding of the ESRE motif, different activated form of the transcription factor (40), or mere non-specific bindings. To increase specificity, we added an additional ESRE sequence into the DNA probe (4XESRE). We observed a decrease of unspecific bindings, with one of the bands (Shift) to remain. This DNA binding activity was enhanced by both phenanthroline and rotenone exposure (**Figure 2B**, **Figure 2-source data 3-4**). This result confirmed that a stress-inducible nuclear factor(s) binds the ESRE motif after mitochondrial damage and suggested that it may be required for the transcription of the ESRE genes. Using the same conditions, bound materials were eluted and subjected to tandem MS/MS analysis for identification of potential peptide fragments.

The gene products identified by mass spectrometry were categorized by using the ‘SRA’ binning system and the ‘iBAQ’ score. The ‘SRA’ or ‘Strict, Relaxed, and All’ binning approach utilizes tiered metrics to score gene identification quality, in which the identified ‘Strict’ genes products pass a 1% FDR cutoff (41). Meanwhile, the ‘iBAQ’ scores were calculated based on peptide peak intensities and number of potential peptides, comparable to the absolute protein quantity. We searched for proteins that passed the ‘SRA’ binning system as ‘Strict’ and are enriched in rotenone-treated samples, as compared to DMSO control, yielding 75 candidates.

To establish a role in ESRE function, each gene predicted to encode one of these proteins was knocked down via RNAi in a strain of *C. elegans* carrying a 3x tandem repeat of the ESRE consensus sequence driving a GFP reporter (3xESRE::GFP) (21). Activation of the reporter was induced using 50 μM rotenone. Amongst the candidate genes, only RNAi targeting *fib-1/Fibrillarin* and *nol-56/Nop56* reduced reporter expression (**Figure 3**). After identifying a role for FIB-1 and NOL-56, we also tested *nol-58/Nop58(RNAi)*, which is a third member of the box C/D snoRNP complex (**Figure 1A**) but was not identified in the affinity-purified material. *nol-58/Nop58(RNAi)* also reduced ESRE expression (**Figure 3**). Current understanding is that the assembly of box C/D snoRNPs occurs via FIB-1/Fibrillarin and NOL-58/Nop58 independently binding the snoRNA and then NOL-56/Nop56 associates with the complex but does not bind to the snoRNA alone (42). Since knockdown of any of the genes for these three proteins reduces ESRE signaling, it seems likely that the snoRNP complex as a whole is binding to ESRE.

**Figure 3.**
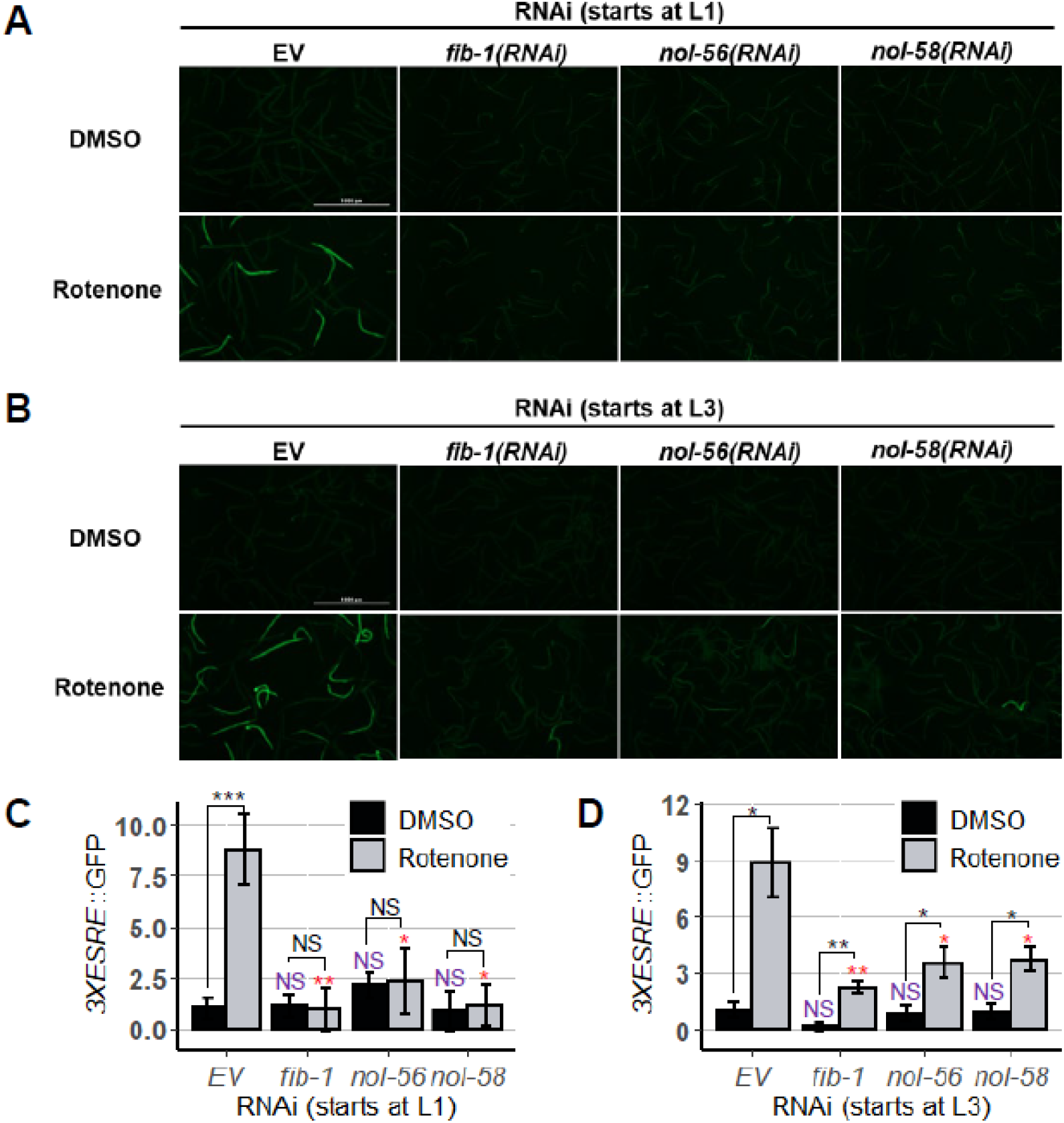
RNAi targeting members of box C/D snoRNP reduced ESRE expression. **(A, B)** Fluorescent images and **(C, D)** quantification of GFP fluorescence of *C. elegans* carrying *3XESRE*::GFP reporters that were reared on *E. coli* expressing empty vector *(EV), fib-1(RNAi), nol-56(RNAi)*, or *nol-58(RNAi)*. Worms were treated for 8 h with vehicle (DMSO) (top) or 50 μM rotenone (bottom). RNAi treatment was started at **(A, C)** L1 or **(B, D)** L3 stage. Representative images are shown; three biological replicates with ~400 worms/replicate were analyzed. Error bars represent SEM. *p* values were determined from one-way ANOVA followed by Dunnett’s test, and Student’s *t*-test. All fold changes were normalized to DMSO-*EV* control. NS not significant, \**p* < 0.05, \*\**p* < 0.01, *** *p* < 0.001.

Although worms reared on RNAi targeting *fib-1/FBL*, *nol-56/Nop56*, or *nol-58/Nop58* exhibited reductions in ESRE signaling, they also showed clear signs of reduced growth and development, resulting in smaller adults, which is consistent with a previous report (43). To avoid this effect, we performed the same experiment, but exposed worms to RNAi starting at the L3 stage instead. Exposure at a later stage of development can circumvent some of the developmental effects, but it can also reduce penetrance (44). However, knockdowns at L3 also reduced ESRE activation upon stress (**Figure 3B, D**). These data confirm that box C/D snoRNPs are required for ESRE pathway gene regulation.

### Box C/D snoRNPs also regulate UPR^mt^ and MAPK^mt^

To assess whether knocking down box C/D snoRNPs only affected the ESRE pathway or impacted other mitochondrial stress responses, we measured the expression of downstream effectors for UPR^mt^ and MAPK^mt^ using GFP-based reporters. Young adult worms carrying *Phsp-6*::GFP (for UPR^mt^) were reared on plates containing all pairwise mixtures of RNAi: empty vector or *spg-7/SPG7(RNAi)* with empty vector, *fib-1(RNAi)*, *nol-56(RNAi)* or *nol-58(RNAi)*. SPG-7/SPG7 is a mitochondria-resident protease that is required for normal organellar function (45); *spg-7(RNAi)* efficiently induces UPR^mt^ (18, 46, 47). As with the ESRE pathway, knockdown of *fib-1*, *nol-56*, or *nol-58* reduced the ability of the UPR^mt^ to respond to stress, regardless of whether RNAi was begun at the L1 or L3 stage (**Figure 4A, B**). Importantly, we also observed significant reduction of the basal level of reporter gene expression. Unlike induced ESRE function following L1 RNAi, however, neither condition completely disrupted UPR^mt^ activity (compare **Figure 3** and **Figure 4A**).

**Figure 4.**
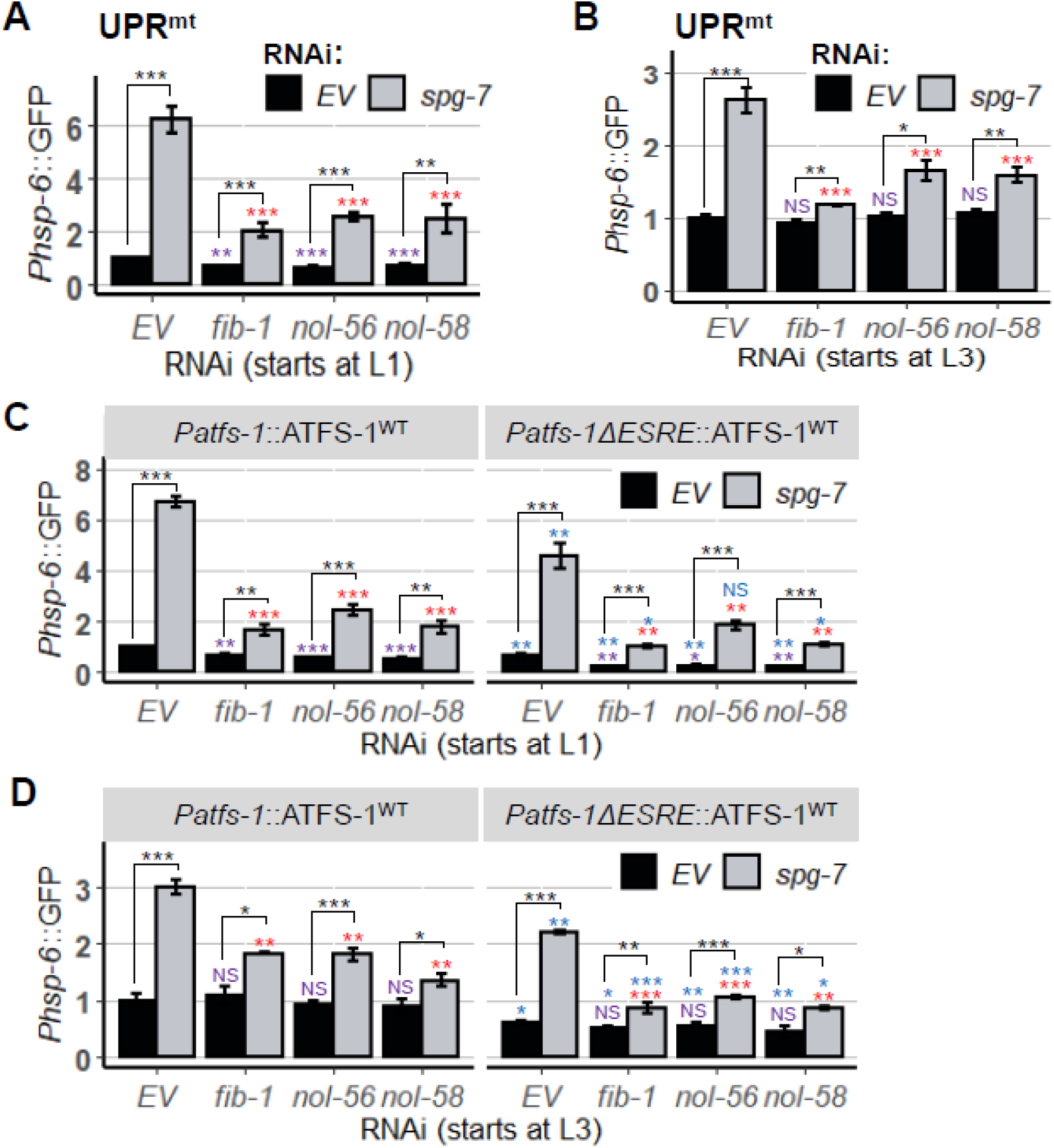
*Box C/D snoRNPs(RNAi)* reduced UPR^mt^ expression. Quantification of GFP fluorescence of *C. elegans* carrying *Phsp-6*::GFP reporter that were reared on *E. coli* expressing empty vector *(EV)* or RNAi targeting box C/D snoRNP members: *fib-1/FBL, nol-56/Nop56*, and *nol-58/Nop58*. RNAi was doubled with empty vector *(EV)* or *spg-7(RNAi)*. In **(C, D)**, *Phsp-6*::GFP reporter strains were wild type or crossed with *Palfs-1ΔESRE*::ATFS-1^WT^. Three biological replicates with ~400 worms/replicate were analyzed. Error bars represent SEM. *p* values were determined from one-way ANOVA, followed by Dunnett’s test, and Student’s *t*-test. All fold changes were normalized to *EV* control. NS not significant, \**p* < 0.05, ** *p* < 0.01, *** *p* < 0.001.

Previously, we demonstrated relationships between the ESRE network and the UPR^mt^ and MAPK^mt^ pathways (21). For example, UPR^mt^ and MAPK^mt^ activity normally places a brake on the ESRE network by limiting the production of ROS that activate ESRE. We also identified an ESRE motif in the promoter region of ATFS-1/ATF5 that was required for its full expression.

Using *spg-7(RNAi)* to induce UPR^mt^, we compared *Phsp-6*::GFP reporter expression in strains with or without the ESRE motif in *atfs-1* promoter on vector control or after disrupting the box C/D snoRNP complex. As expected, knocking down the protease induced GFP expression in each condition. Similar to our previous findings, basal and induced expression of *Phsp-6*::GFP was lower when ESRE motif was removed (blue significance marks in **Figures 4C-D**). Adding RNAi targeting the box C/D machinery to the ESRE deletion changed basal expression only when RNAi was initiated at L1 stage. Induction of *hsp-6::GFP* by *spg-7(RNAi)* was lower when compared to empty vector or when compared to induced conditions with an intact promoter and box C/D RNAi. These data indicate that the box C/D complex regulates UPR^mt^ both via modulation of ESRE pathway (due to a presence of ESRE motif, which is required for full expression of *atfs-1*) and independently of ESRE.

Contrary to what was observed for UPR^mt^, *fib-1(RNAi)*, *nol-56(RNAi)*, and *nol-58(RNAi)* caused a statistically significant increase in basal expression level of the *Ptbb-6*::GFP MAPK^mt^ reporter (**Figure 5A**), indicating that the box C/D snoRNP complex is directly or indirectly involved in repressing basal expression of the MAPK^mt^ pathway. This difference disappeared if RNAi targeting the box C/D snoRNP complex was initiated at the L3 stage (**Figure 5B**). RNAi at either stage had no effect on *Ptbb-6*::GFP expression after induction via *spg-7(RNAi)*. As expected, *Ptbb-6*::GFP expression was at least partially dependent upon PMK-3/MAPK14, both under wild-type and box C/D snoRNP conditions (**Figure 5C**). Previously we demonstrated that ATFS-1 plays a role in repressing *tbb-6* expression (21). Since we showed above that ATFS-1 activity can depend on box C/D snoRNPs, we tested whether *fib-1(RNAi)*, *nol-56(RNAi)*, or *nol-58(RNAi)* would affect ATFS-1-mediated repression *tbb-6*. In each case, *atfs-1* knockdown was indistinguishable from *atfs-1; snoRNPs* double RNAi. (**Figure 5D**). Combined, these data argue that the box C/D complex regulates basal expression of the MAPK^mt^ stress response system via altering levels of ATFS-1. We also speculate that the absence of changes for basal ESRE reporter expression are due to consistent observations that ESRE network activation must be spurred by recognition of stress, while *Phsp-6::*GFP and *Ptbb-6::*GFP exhibit low levels of expression even in the absence of stress (18).

**Figure 5.**
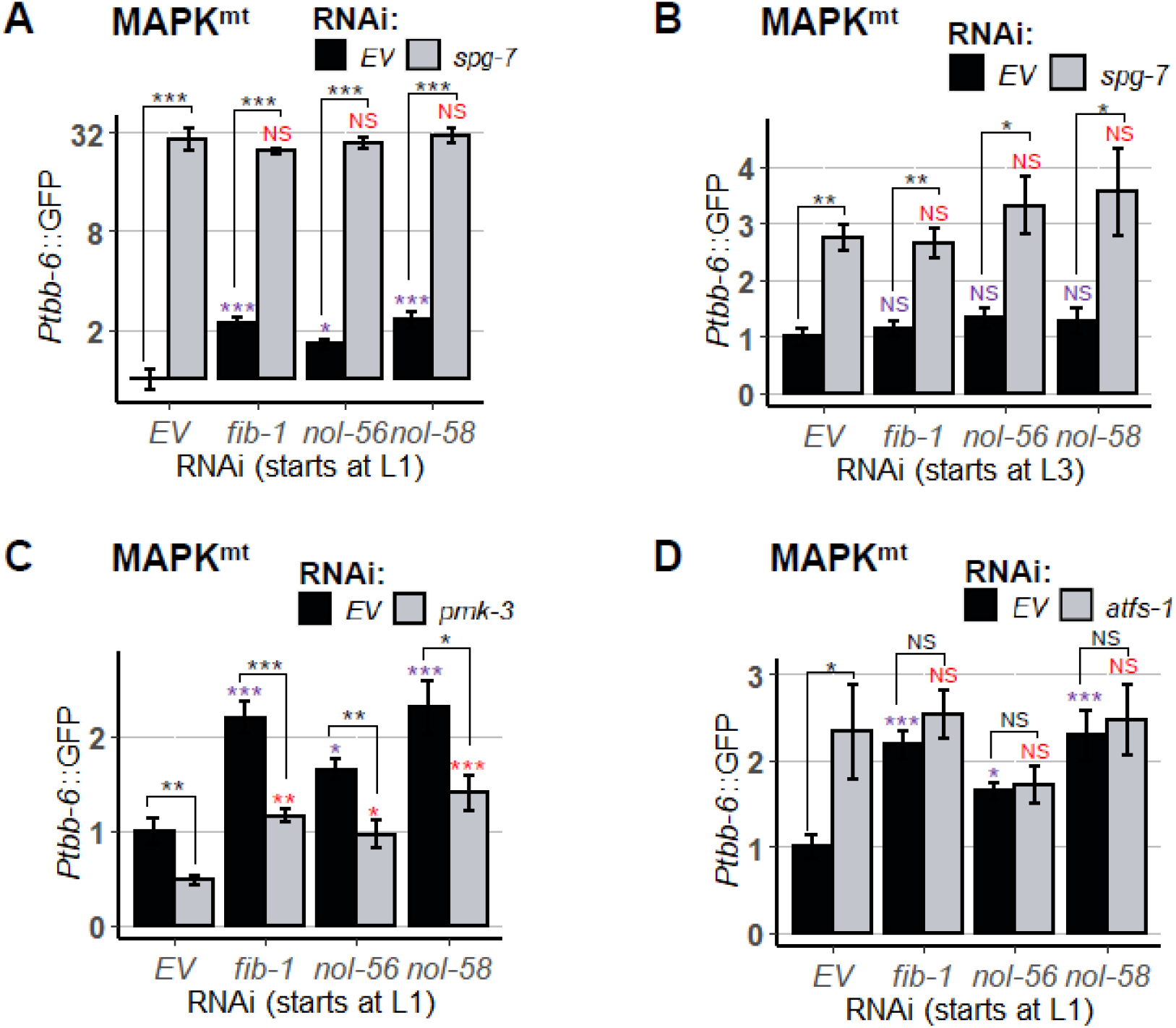
The loss of box C/D snoRNPs increased MAPK^mt^ expression. Quantification of GFP fluorescence of *C. elegans* carrying *Ptbb-6*::GFP reporter that were reared on *E. coli* expressing empty vector *(EV)* or RNAi targeting box C/D snoRNP members: *fib-1/FBL, nol-56/Nop56*, and *nol-58/Nop58*. RNAi was doubled with empty vector *(EV)* or **(A-B)** *spg-7(RNAi)*, **(C)** *pmk-3(RNAi)*, or **(D)** *atfs-1(RNAi)*. Three biological replicates with ~400 worms/replicate were analyzed. Error bars represent SEM. *p* values were detennined from oneway ANOVA, followed by Dunnett’s test, and Student’s *t*-test. All fold changes were normalized to *EV* control. NS not significant, \**p* < 0.05, ** *p* < 0.01, *** *p* < 0.001.

We further asked whether localization of these snoRNPs in the nucleolus is necessary for the regulation to take an effect. We knocked down *ruvb-1/RUVB*, an AAA+ ATPase that promotes box C/D snoRNPs assembly and localization to nucleoli (43). Worms reared on *ruvb-1(RNAi)*-expressing *E. coli* at L1 did not show growth arrest. *ruvb-1/RUVB* knockdown markedly reduced ESRE expression following stress (**Supplementary Figure 1A**). However, induced expression of UPR^mt^ was not affected (**Supplementary Figure 1B**) and increased in basal expression of MAPK^mt^ reporter was also more modest than what was observed for C/D snoRNPs knockdown (**Supplementary Figure 1C**). This suggests that localization of box C/D snoRNPs may affect some but not all of the responses of mitochondrial surveillance.

### Disruption of box H/ACA snoRNP machinery does not affect mitochondrial surveillance pathways

One possible explanation for the phenomena that we observed was that the reduction of 2’-O-methylation of rRNA compromised normal ribosomal function. If this were true, other broad-scale ribosomal changes should have similar outcomes. As previously noted, the conversion of dozens to hundreds of uridine residues to pseudouridine in rRNA is catalyzed by box H/ACA snoRNPs (48, 49). RNAi was used to target the genes encoding three of the four essential proteins for box H/ACA complex activity: *nola-3/Nop10, Y48A6B.3/Nhp2, and Y66H1A.4/Gar1*, and basal and induced expression of mitochondrial stress reporters were assessed. We observed no significant change of expression for any of the mitochondrial surveillance pathways tested in either basal or induced conditions (**Supplementary Figure 2**). This result indicates that the function of box C/D snoRNPs in regulating mitochondrial homeostasis is specific.

### Suppressing translation does not recapitulate changes in mitochondrial surveillance caused by disrupting the box C/D snoRNP complex

2’-O-methylation has a number of effects on ribosome maturation and stability. One possible consequence of disrupting ribosomal biology is a global reduction in translation (50, 51). Translation efficiency is known to be a target of surveillance in *C. elegans* (1, 2). Importantly it was recently shown that *fib-1/FBL* knockdown activates *irg-1*, an innate immune reporter, (38) and we recapitulated these data and observed increased reporter expression on *nol-56(RNAi)* and *nol-58(RNAi)* (**Supplementary Figure 3A)**. *irg-1* is known to respond to translational inhibition, including exposure to exotoxin A, hygromycin (1, 2), or cycloheximide (**Supplementary Figure 3B**). For these reasons, Tiku *et al*, hypothesized that knockdown of fibrillarin results in the suppression of translation, triggering activation of innate immunity. Thus, we set out to explore whether the decreased levels of the ESRE reporter following *fib-1(RNAi)* are caused by the same mechanism. We tested whether snoRNPs regulate ESRE by modulating translation. We used RNAi to target the genes encoding components of the eukaryotic 48S transcription initiation complex, including *clu-1/eIF3A, inf-1/eIF4A, ife-2/eIF4E, ifg-1/eIF4G*, and *T12D8.2/eIF4H*. Since RNAi targeting *ifg-1/eIF4G* and *inf-1/eIF4A* compromised development when RNAi was started at the L1 stage, subsequent experiments were performed by feeding RNAi starting at the L3 stage of development.

Disrupting the 48S complex had no consistent effect on the expression of reporters for the mitochondrial surveillance pathway, with the exception of *inf-1/eIF4A(RNAi)*, which showed a reduction in rotenone-mediated ESRE activation and *spg-7*-mediated UPR^mt^ activation, and increased basal expression of MAPK^mt^ (**Figure 6**). However, induction of the *Phsp-6::GFP* reporter by *spg-7(RNAi)* was also significantly decreased for *clu-1/eIF3A(RNAi), ifg-1/eIF4G(RNAi), and ife-2/eIF4E(RNAi)*, suggesting some specialized interactions (**Figure 6B**).

**Figure 6.**
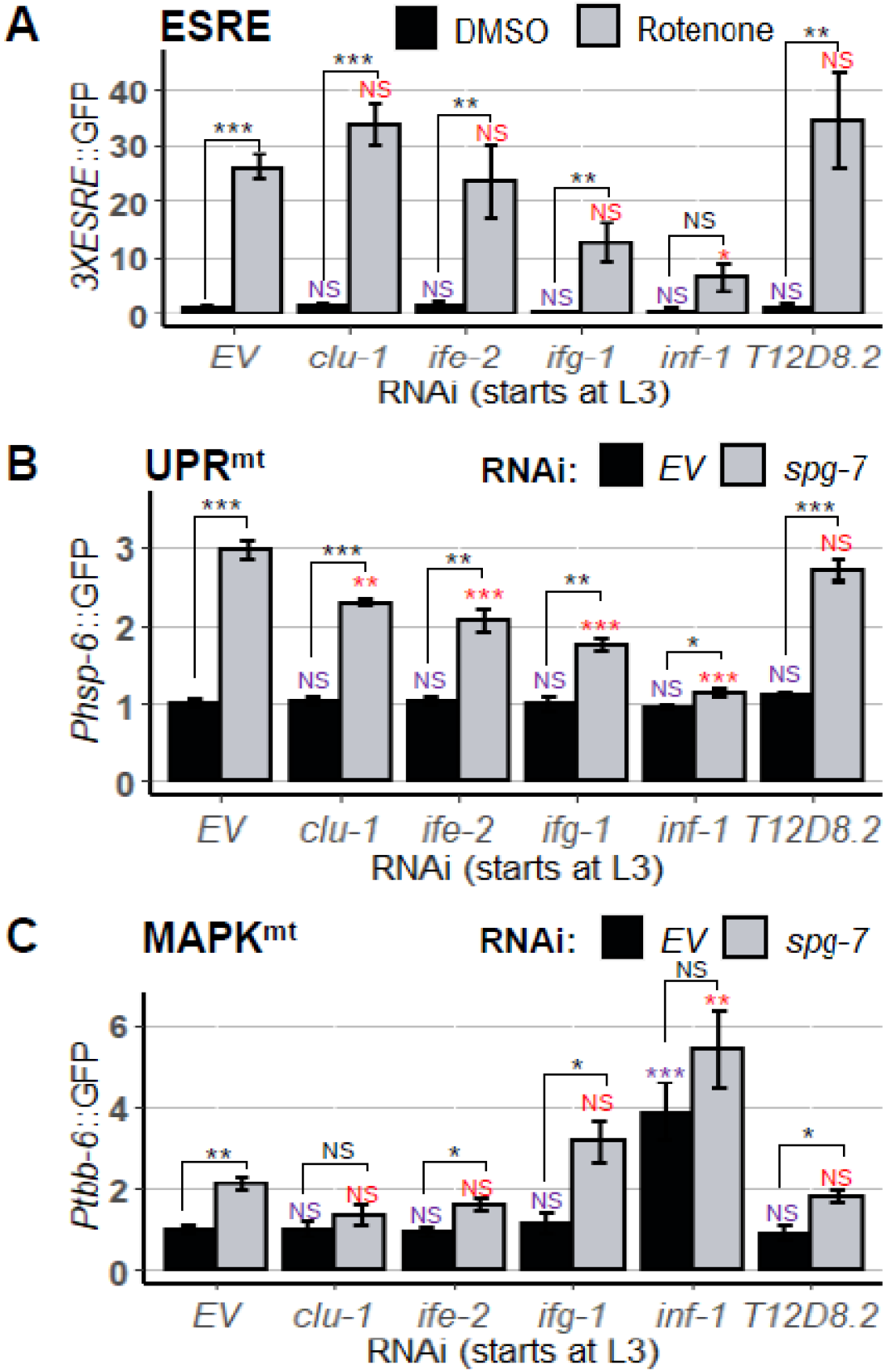
RNAi targeting eukaryotic initiation factors partially affected the mitochondrial surveillance pathways. Quantification of GFP fluorescence of *C. elegans* carrying **(A)** *3XESRE*::GFP, **(B)** *Phsp-6*::GFP, and **(C)** *Ptbb-6*::GFP reporters that were reared on *E. coli* expressing empty vector *(EV)* or RNAi targeting several euKaryotic initiation factors: *clu-1/elF3A, ife-2/elF4E, ifg-1/elF4G, inf-1/elF4A*, and *T1208.2/elF4H*. In **(A)**, worms were treated for 8 h with vehicle (DMSO) or 50 μM rotenone. In **(B, C)**, double RNAi was performed with empty vector *(EV)* or *spg-7(RNAi)*. Three biological replicates with ~400 worms/replicate were analyzed. Error bars represent SEM. *p* values were determined from one-way ANOVA, followed by Dunnett’s test, and Student’s *t*-test. All fold changes were normalized to DMSO-EVor *EV* control. NS not significant, \**p* < 0.05, ** *p* < 0.01. *** *p* < 0.001.

As a final test, we treated reporter worms for each of the three mitochondrial surveillance pathways with the chemical translational inhibitor cycloheximide under conditions where *irg-1* activation was observed (see **Supplementary Figure 3**). Cycloheximide did not alter ESRE or MAPK^mt^ reporter expression (**Supplementary Figure 4**). These results indicated that general translational reduction is unlikely to be the mechanism that underlies box C/D snoRNP regulation of the ESRE mitochondrial surveillance network.

### Box C/D snoRNPs repress innate immune responses

As mentioned above, fibrillarin knockdown has previously been linked to increased pathogen resistance (38), and we and others observed increased expression *Pirg-1::GFP*, an immune reporter, in *fib-1(RNAi)* worms (**Supplementary Figure 3**). To test whether disruption of the box C/D snoRNP complex induced other cellular defense pathways, we monitored the expression of *Pirg-5*::GFP reporter, which is activated by a number pathogens and xenobiotics (52, 53) and is relatively insensitive to translational inhibition (**Figure 7A** and (2)).

**Figure 7.**
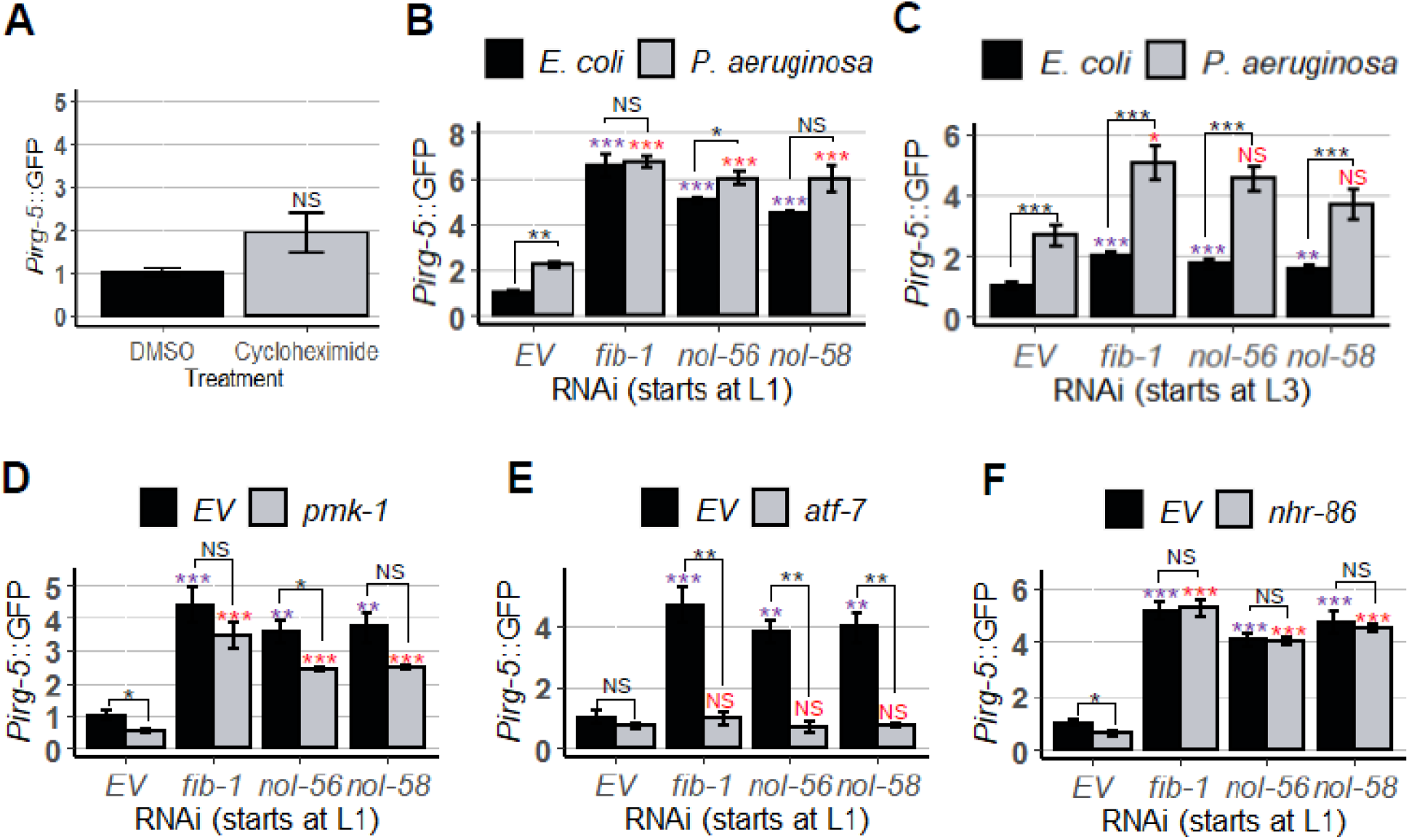
Knockdown of box C/D snoRNPs increased immune response. Quantification of GFP fluorescence of *C. elegans* carrying *Pirg-5*::GFP reporter. In **(A)**, worms were treated for 8 h with vehicle (DMSO) or translation elongation inhibitor cycloheximide. In **(B-F)**, worms were reared on *E. coli* expressing empty vector *(EV)* as control and *fib-1(RNAi), nol-56(RNAi)*, or *nol-58(RNAi)*. RNAi treatment was started at **(B, D, E, F)** L1 or **(C)** L3 stage. In **(B-C)**, young adult worms were transferred onto plates containing *E. coli* or *P. aeruginosa* and let roamed for 8 h. In **(D-F)**, GFP fluorescence were measured on basal levels. Three biological replicates with ~400 worms/replicate were analyzed. Error bars represent SEM. *p* values were determined from one-way ANOVA, followed by Dunnett’s test, and Student’s *t*-test. All fold changes were normalized to *EV* control on *E. coli*. NS not significant, \**p* < 0.05, ** *p* < 0.01, *** *p* < 0.001.

Worms carrying the *Pirg-5*::GFP reporter were reared on either empty vector or RNAi targeting components of the box C/D snoRNP complex starting at L1, and then GFP expression was evaluated in young adults. Interestingly, this disruption activated *irg-5* more strongly than *P. aeruginosa* infection on agar in empty vector controls (**Figure 7B**). It is also worth noting that *P. aeruginosa* infection of worms with RNAi targeting *fib-1* and *nol-58* did not increase GFP expression any further than in their uninfected counterparts, suggesting that *irg-5* induction may have already been maximized.

As we had observed previously, box C/D RNAi affected ESRE induction differently when RNAi was started at L1 vs L3. Specifically, increased basal induction was much stronger when RNAi feeding was initiated at the L1 stage (**Figure 7B-C**). This observation is also consistent with our interpretation that *Pirg-5::GFP* expression is nearly saturated when box C/D is knocked down early in development (**Figure 7B**). These results indicate that early developmental C/D snoRNP complexes are required for appropriate innate immune function later in development.

As *irg-5* was not significantly upregulated by the presence of cycloheximide, i.e., by translational repression, snoRNPs are likely to affect innate immune pathways via multiple mechanisms. *irg-5* is known to be controlled by several transcriptional regulators, including PMK-1/p38 MAPK and ATF-7/ATF7 (53), both which are established regulators of innate immunity in *C. elegans* (54). ATF-7/ATF7 functions downstream of PMK-1/p38 MAPK, but it can regulate *irg-5* activation in response to small molecule immune stimulator RPW-24 independently of PMK-1/p38 MAPK (53). NHR-86/HNF4 also regulates *irg-5* expression, specifically in response to the xenobiotic compound RPW-24 (55). To determine whether any of these transcriptional regulators were involved in box C/D regulation of innate immunity, we compared *Pirg-5::GFP* expression in worms with double RNAi targeting *fib-1/Fibrillarin, nol-56/Nop56*, or *nol-58/Nop58* and *pmk-1(RNAi)*, *atf-1(RNAi)*, or *nhr-86(RNAi)*.

Both *pmk-1(RNAi)* (**Figure 7D**) and *atf-7(RNAi)* (**Figure 7E**) reduced *Pirg-5::GFP* expression, with the latter virtually abolishing its expression. *nhr-86(RNAi)* (**Figure 7F**) had no apparent effect. These data suggest that the effects of the box C/D snoRNP complex on *irg-5* are upstream of its known regulation by the ATF-7/ATF7.

This led us to question whether ATF-7 is involved in the regulation of other reporters that are differentially expressed under box C/D disruption. To test the *3XESRE::GFP* reporter, worms were reared on combinations of empty vector, *fib-1(RNAi)*, *atf-7(RNAi)*, or both starting at the L3 stage. Young adults were then exposed to rotenone and reporter induction was observed. Knockdown of *atf-7* was indistinguishable from vector control (red “NS” mark), and *atf-7(RNAi);fib-1(RNAi)* was not different from just *fib-1(RNAi)* (blue “NS” mark) (**Figure 8A**). This indicates that the box C/D regulation of ESRE is independent of ATF-7 and is different from the regulation of *irg-5*.

**Figure 8.**
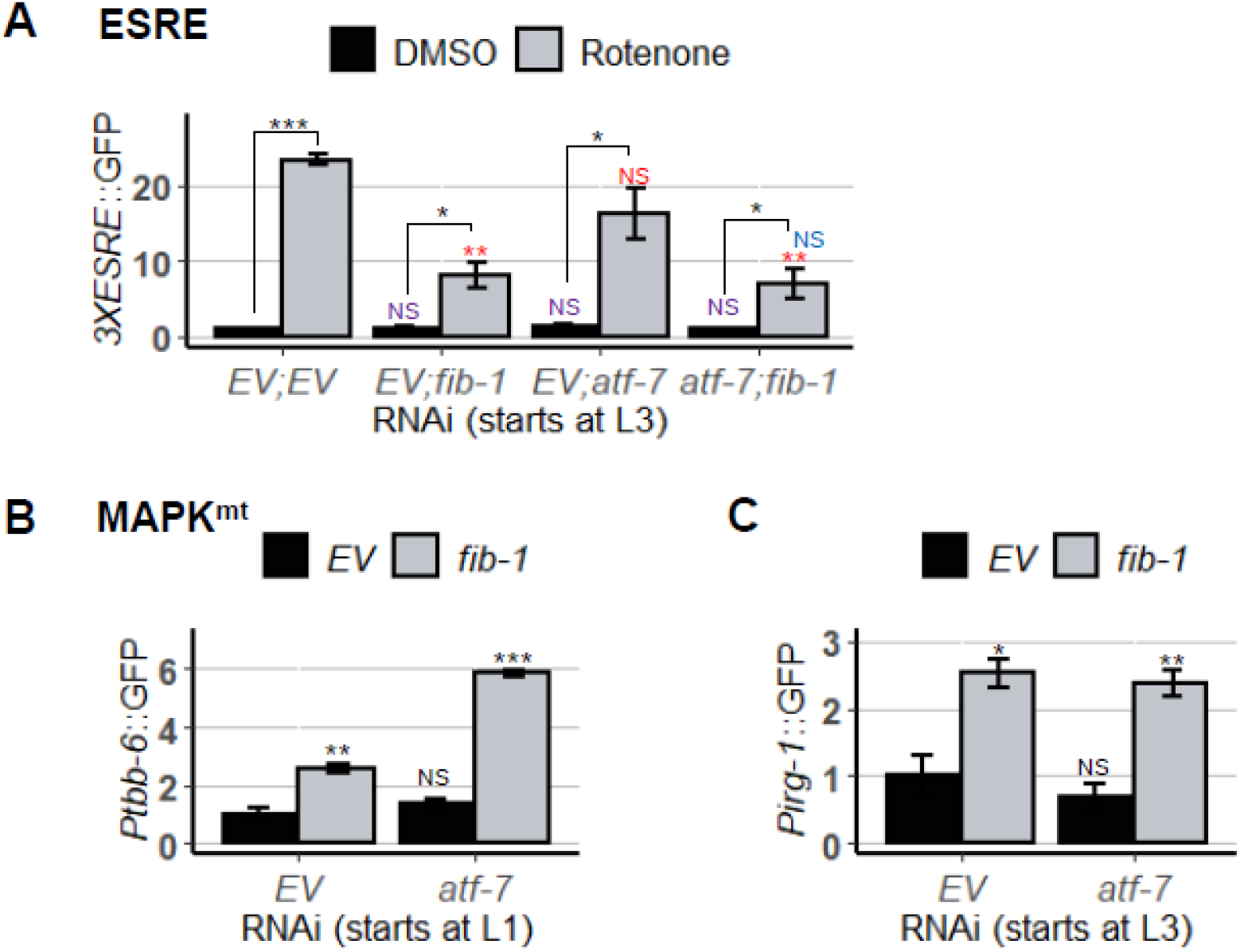
Knockdown of *atf-7* did not affect mitochondrial surveillance pathways. Quantification of GFP fluorescence of *C. elegans* carrying **(A)** *3XESRE*::GFP, **(B)** *Ptbb-6*:GFP, or **(C)** *Pirg-1*::GFP reporters. Worms were reared on *E. coli* expressing empty vector *(EV)* as control or RNAi targeting *fib-1, atf-7*, or *atf-7* and *fib-1*. Three biological replicates with ~400 worms/replicate were analyzed. Error bars represent SEM. *p* values were determined from one-way A NOVA, followed by Dunnett’s test, and Student’s *t*-test. All fold changes were normalized to *EV* control. NS not significant, \**p* < 0.05, ** *p* < 0.01, *** *p* < 0.001.

We also tested whether the increase in basal expression of *Ptbb-6::GFP* after *fib-1(RNAi)* was related to ATF-7 by rearing the reporter on *fib-1(RNAi)* alone or with *atf-7(RNAi)* starting at the L1 larval stage. As expected, basal levels of the reporter were increased after *fib-1(RNAi)* but not by *atf-7(RNAi)* alone (**Figure 8B**). However, we did observe an additive effect on *Ptbb-6::GFP* in *atf-7(RNAi);fib-1(RNAi)*, suggesting that the two genes work together to limit inappropriate expression of the MAPK^mt^ pathway.

*atf-7(RNAi)* did not affect expression of *irg-1*, an immune effector whose basal expression is also upregulated upon box C/D snoRNP knockdown (**Figure 8C**). This suggest that box C/D snoRNPs exhibit complex modulation of innate immune pathways. Although our previous experiments had ruled out a role box H/ACA snoRNPs, in the regulation of mitochondrial surveillance (**Supplementary Figure 2**), we wanted to investigate their role in innate immunity. A knockdown of box H/ACA did not invoke any response from the immune genes *irg-1* and *irg-5* (**Supplementary Figure 5**), confirming the specificity of box C/D snoRNPs.

### The loss of box C/D snoRNPs reduced survival in liquid-based P. aeruginosa killing assay

To test whether the loss of box C/D snoRNPs had physiologically relevant consequences, we performed *P. aeruginosa* Liquid Killing and Slow Killing assays to measure survival. Although both assays use the same pathogen, the virulence and pathogenic mechanisms and the host defenses differ. In the Liquid Killing assay, host death occurs due to the production of the siderophore pyoverdine, which is secreted by the bacterium to obtain iron (26, 56). Pyoverdine enters host tissue and removes iron from mitochondria, causing sufficient damage to inflict death (57, 58). This damage also activates the ESRE mitochondrial surveillance network, which is important for host defense (20).

The Slow Killing assay is more traditional form of bacterial pathogenesis, where the host intestine is colonized by the pathogen, and killing involves quorum sensing, although the precise cause of death has not yet been determined (59, 60). Slow Killing activates the conventional NSY-1/SEK-1/PMK-1 MAPK pathway, which is the most common antibacterial defense in *C. elegans* (61–63). Interestingly, there appears to be little overlap between the two defense networks, as the Slow Killing pathway has no activation of ESRE and PMK-1 activity is actually detrimental for survival under Liquid Killing conditions (25, 64).

Worms were reared on RNAi targeting *fib-1/Fibrillarin*, *nol-56/Nop56*, or *nol-58/Nop58* from the L3 larval stage, and then young adults were exposed to *P. aeruginosa* strain PA14 either under Liquid Killing or Slow Killing conditions. As anticipated based on the role of ESRE in improving survival in Liquid Killing and the observation that box C/D snoRNP knockdown compromises ESRE function, removal of box C/D function strongly reduced host survival (**Figure 9A**). Interestingly, we saw the opposite in Slow Killing, where box C/D snoRNP RNAi slightly, but statistically significantly, increased host survival during *P. aeruginosa* intestinal infection (**Figure 9B**). This is consistent with a prior report that FIB-1 reduces host survival during infection (38).

**Figure 9.**
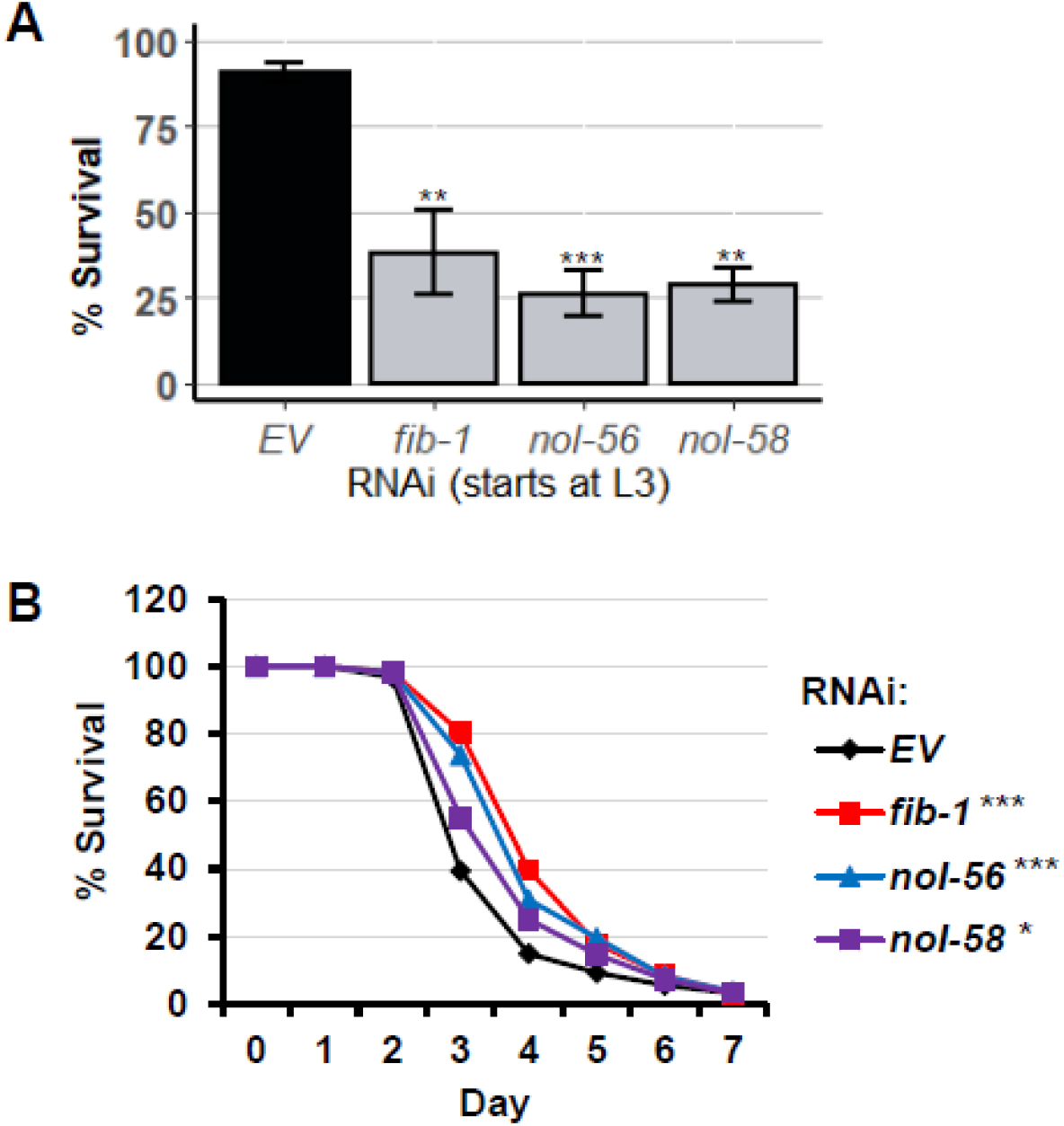
The loss of box C/D snoRNPs had opposing effect on two *P. aeruginosa* pathogenesis assays. Survival of *glp-4(bn2)* worms grown on RNAi strains targeting box C/D snoRNPs in **(A)** Liquid Killing and **(B)** Slow killing assays. Three biological replicates with ~400 worms/replicate for LK or ~150 worms/replicate for SK were analyzed. The average of all replicates is shown for each panel. Error bars represent SEM. Fold changes were normalized to *EV* control. *p* values were determined from one-way ANOVA. followed by Dunnett’s test for LK or log-rank test for SK. \**p* < 0.05, ** *p* < 0.01, *** *p* < 0.001.

## Discussion

In this study, we identified a role for the box C/D snoRNP complex in regulating the switch between mitochondrial surveillance and innate immunity. Using biochemical approaches, we made the unexpected discovery that FIB-1/Fibrillarin and NOL-56/Nop56 were associated with a tandem repeat of the ESRE motif. This is unexpected for two reasons. First, the RNA sequences that box C/D snoRNAs are named for, the C (**RUGAUGA**) and D (**CUGA**) boxes, have been well-studied. These bear very little sequence similarity to the ESRE motif (**TCTGCGTCTCT**). Although these are each consensus and have some variability, there is essentially no match, so it seems unlikely that the proteins are recognizing the ESRE site directly. Second, as noted above, NOL-56/Nop56 appears to recognize one of the protein components of box C/D snoRNPs, which means that the snoRNA is likely to be present.

Box C/D snoRNPs have recently been linked to an increasing variety of function, including rare cases of guiding RNA editing (65), tRNA methylation (66), and even association with mRNA (including an ‘orphan’ snoRNA with no known rRNA target that destabilizes several mRNAs) (67, 68). Additionally, snoRNAs are more frequently found in the cytoplasm after exposure to oxidative stress or heat shock (69–71), suggesting the possibility that snoRNAs could be leaving the nucleolus to regulate ESRE genes by methylating mRNAs. But this also seems to be an unlikely mechanism for what we observed. Approximately 50 box C/D snoRNAs are predicted in the genome of *C. elegans*. In contrast, the ESRE nucleotide motif is present in the promoter region of ~8% of predicted genes (22). This numerical discrepancy makes it rather unlikely that a single snoRNA is responsible for the regulation of all of them, unless it recognizes the ESRE consensus sequence. None of the snoRNAs predicted in the *C. elegans* genome are obvious candidates for recognizing the ESRE motif; in all cases we are aware of, the hybridization sequence is located between the C and D boxes of the snoRNA, and none of these matched ESRE.

An alternative hypothesis is that stress signals change snoRNP expression or targeting, resulting in changes to rRNA modification patterns. Most known box C/D snoRNP targets are in rRNA, and it is worth noting that some of these are not saturated and the degree to which some sites are modified is associated with cellular stress (72). This has led to the suggestion that there may be different populations of ribosomes within cells, some with specialized functions. This could include ribosomes that more efficiently express stress-responsive genes, altering the transcriptome of the cells. The well-known integrated stress response, where eIF-2α is phosphorylated and cap-dependent mRNA translation is substantially decreased in favor of translation from structured mRNA elements called internal ribosomal entry sites, or IRESes (16), is an example of one such condition. Changes to the ribosome could then facilitate specification for structured mRNAs.

In this case, we would see increased ESRE gene expression during stress. While we do see increased expression of ESRE genes during stress, this is at least partially due to transcriptional changes, and we have not yet seen evidence of translational differences. However, ESRE genes are not known to be activated by translational inhibitors like *P. aeruginosa* exotoxin A or hygromycin (1). Targeting the 48S pre-initiation complex here (**Figure 6A**) also did not activate ESRE gene regulation. Additionally, we saw no changes in ESRE gene expression when components of the box H/ACA snoRNP complex were disrupted. rRNA modification by these ribonucleoprotein complexes are also important and would also be expected to affect transcription if this were mechanism of ESRE gene regulation. Additionally, stress-responsive ribosomal modifications do not explain the association of the box C/D snoRNP complexes with the ESRE motif.

Importantly, we found that two immune effectors, *irg-1* (**Supplementary Figure 3** and (38)) and *irg-5*, were upregulated by the absence of box C/D snoRNPs. This is consistent with many reports that disruption of core cellular processes activates innate immune processes (1, 9, 29). In this case, the loss of box C/D snoRNPs activated *irg-1* via translation suppression (38) and *irg-5* through an unknown mechanism. Interestingly, only the loss of ATF-7/ATF7, a transcription factor that regulates *irg-5* response to pathogen attack (and partially to xenobiotic compound), was able to completely abolish *irg-5* induction by RNAi targeting box C/D snoRNP machinery. This indicated that ATF-7/ATF7, partially independently of PMK-1/p38 MAPK, regulated *irg-5* expression in response to the loss of box C/D snoRNPs. This is similar to *irg-5* induction by the immune stimulator RPW-24 (53). However in this case, knockdown of *nhr-86/HNF4* did not abolish *irg-5* expression, suggesting a different biological significance of the activation of this ATF-7/ATF7-dependent immune pathway.

We propose that the box C/D snoRNPs act as a molecular switch that activate quality control pathways while inhibiting immune responses (**Figure 10**). The purpose of this novel mechanism may be to allow mitochondria an opportunity to repair before other cellular defenses (that require extensive energy expenditure) are activated. Future work will focus on understanding the relationships between these systems.

**Figure 10.**
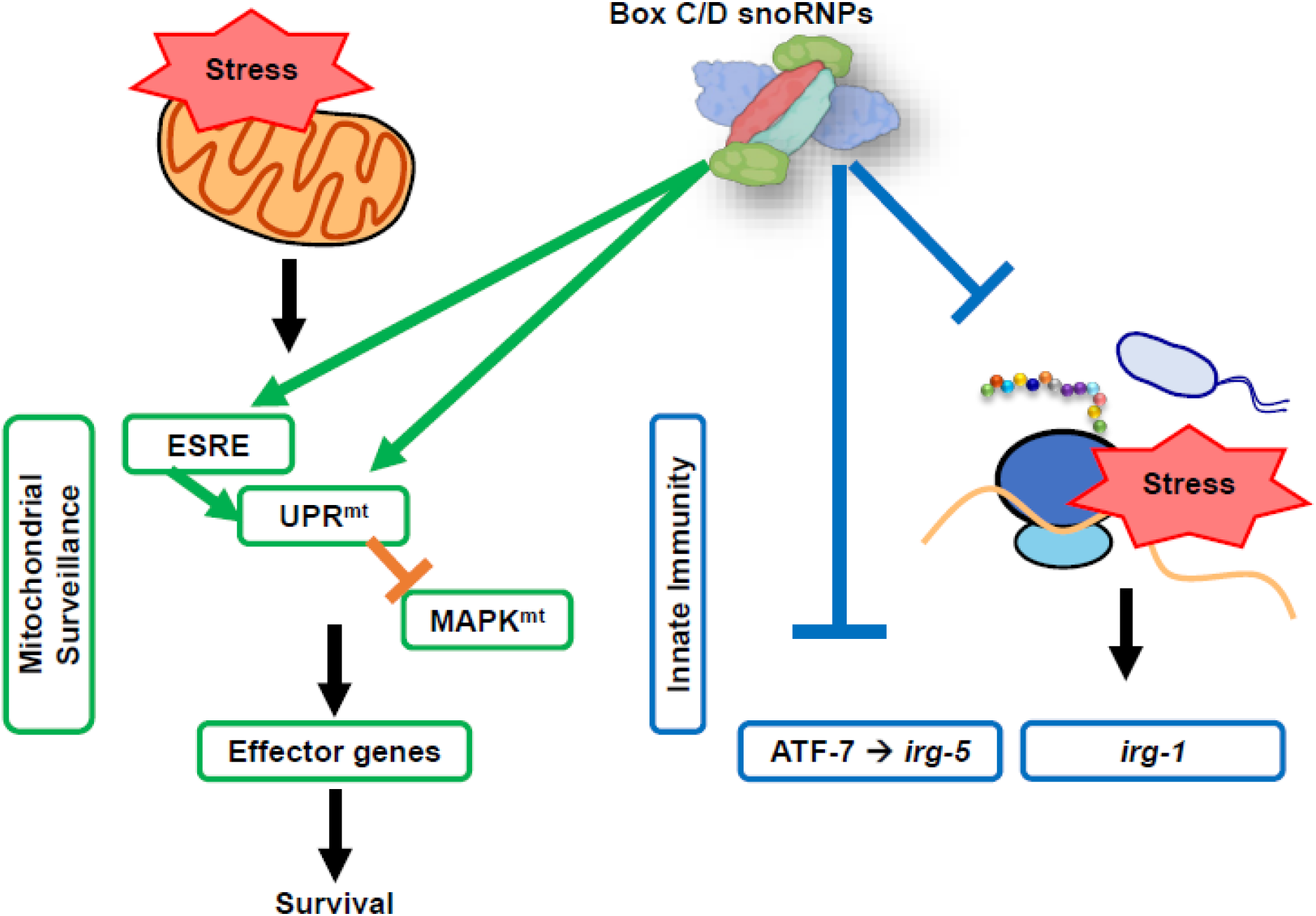
Proposed model of box C/D snoRNPs roles in cellular pathways regulation. Box C/D snoRNPs act as a molecular switch that suppresses innate immunity and activates mitochondrial pathways upon stress.

## Methods

### *C. elegans* strains and maintenance

All *C. elegans* strains were maintained on standard nematode growth medium (NGM) (73) seeded with *Escherichia coli* strain OP50 as a food source and were maintained at 20°C (73), unless otherwise noted. *C. elegans* strains used in this study included N2 Bristol (wild-type), SS104 [*glp-4*(*bn2*)], WY703 |*fdIs2* [*3XESRE*::GFP]; *pFF4*[*rol-6*(*su1006*)]| (27), SJ4100 |*zcIs13* [*Phsp-6*::GFP]|,|*atfs-1*(*et15*); *zcIs9* [*Phsp-60::GFP*]| (74), NVK235 (*zcIs13*; *Patfs-1ΔESRE*::ATFS-1^WT^) (21), SLR115 |*dvIs67* [*Ptbb-6*::GFP + *Pmyo-3*::dsRed]| (18), AY101 |*acIs101* [*pDB09.1*(*Pirg-5*::GFP); *pRF4*[*rol-6*(*su1006*)]]| (52), and AU133 |*agIs17*[*Pmyo-2*::mCherry + *Pirg-1*::GFP]| (2).

Worms were synchronized by hypochlorite isolation of eggs from gravid adults, followed by hatching of eggs in S Basal. 6000 synchronized L1 larvae were transferred onto 10 cm NGM plates seeded with OP50. After transfer, worms were grown at 20°C for 50 hours prior to experiments, or for three days for the next eggs isolation. Young adult worms were used for all assays unless otherwise noted.

### Bacterial strains

RNAi experiments in this study were done using RNAi-competent HT115 obtained from the Ahringer or Vidal RNAi library (75, 76) and were sequenced prior to use. For *P. aeruginosa*, PA14 strain was used (77).

### RNA interference protocol

RNAi-expressing bacteria were cultured and seeded onto NGM plates supplemented with 25 μg/mL carbenicillin and 1 mM IPTG. When double RNAi was performed, bacteria cultures were mixed with a 1:1 ratio. For RNAi experiment starting at L1, 2000 synchronized L1 larvae were transferred onto 6 cm RNAi plates and grown at 20°C for 50 hours prior to imaging or exposure to chemical compounds or pathogens. For RNAi experiment starting at L3, 2500 synchronized L1 larvae were transferred onto 6 cm regular NGM plates seeded with OP50 and grown at 20°C for 20 hours until reaching the L3 stage. Worms were then washed off plates, rinsed three times, and transferred onto RNAi plates. Worms were grown at 20°C for 30 hours on RNAi plates prior to use for experiments.

### Electrophoretic Mobility Shift Assay (EMSA) and Oligo Pull-down

Cytoplasmic and nuclear protein extraction was performed with Pierce Cytoplasmic and Nuclear Extraction Kit according to the manufacturer’s protocol.

EMSA was performed by using LightShift^®^ Chemiluminescent EMSA Kit (ThermoFisher). In short, synchronized young adult *glp-4(bn2)* worms were exposed to heat shock for 16 h, 1,1-phenanthroline 1 mM for 20 h, rotenone 50 μM for 14 h, or DMSO (solvent control). Worms’ nuclear-enriched extract was incubated for 20 minutes at room temperature with biotinylated oligos (3XESRE or 4XESRE) as bait in the appropriate binding conditions (50 ng/μL poly (dI•dC), 5% glycerol, 0.1% NP-40, 2.5 mM MgCl2, 1 mM EDTA, and 20 fmol biotinylated oligos). Binding reactions were then loaded for electrophoresis in a polyacrylamide gel until the dye front had migrated ¾ down the length of the gel. Binding reactions were then transferred to a nylon membrane for 30 minutes at 380 mA. DNA on the membrane was then crosslinked at 120 mJ/cm2 by using a UV-light crosslinking instrument. Biotin-labeled DNA was then detected with a series of detection steps before finally exposed to X-ray film.

Oligo pulldown was performed according to the manual for DynaBeads M-280 Streptavidin (ThermoFisher). In short, magnetic beads were first washed with Binding and Washing buffer (BW 2X) (10 mM Tris-HCl pH 7.5, 1 mM EDTA, and 2 M NaCl). Beads were then coupled with the biotinylated-4XESRE oligos bait for 15 minutes (with BW 1X). Coated beads were resuspended in PBS buffer (0.1 M phosphate, 0.15 M NaCl) pH 7.4 and then incubated with worm nuclear extract for 2 hours at room temperature, washed, and eluted. Elution samples were sent for tandem MS/MS.

### *C. elegans* chemical exposure assays

Synchronized young adult worms were washed from NGM plates seeded with OP50 into a 15 mL conical tube and rinsed three times. Worms were then sorted into a 96-well plate (~100 worms/well). S Basal supplemented with 50 μM rotenone (Sigma), 2 mg/mL cycloheximide (Sigma), or DMSO (solvent control) was then added into the wells of the 96-well plate to a final volume of 100 μL. Worms were imaged with Cytation5 automated microscope every two hours for twenty hours. At least three biological replicates were performed for each experiment.

### *C. elegans* pathogenesis assays

Liquid killing was performed essentially as described (56, 78). 25 synchronized young adult worms were sorted into 384-well plate. Liquid killing medium was mixed with *P. aeruginosa* PA14 (final OD_600_: 0.03), and then added into each well. Plates were incubated at 25°C. At time points, plates were washed three times and worms were stained with SYTOX^TM^ Orange nucleic acid stain for 12 h to stain dead worms. Plates were then washed and imaged for with Cytation5 automated microscope and dead worms were quantified with CellProfiler.

Slow Killing was performed as previously described (79). 50 young adult worms were transferred onto PA14-SK plates and incubated at 25°C. Worms were scored every day for survival curve; dead worms were removed from assay plates.

*P. aeruginosa* exposure to worms carrying *Pirg-5*::GFP (AY101) was performed similarly as SK assay. 500 worms were transferred onto PA14-SK plates. After 8 h, worms were washed off plates into a 96-well plate and washed several times to remove bacteria. Imaging and GFP quantification were performed with Cytation5 automated microscope and Gen5 3.0 software.

### Imaging and Fluorescence Quantification

For visualization of the worm reporter strains AU133, AY101, NVK235, SJ4100, SLR115, and WY703 in 96-well plates, Cytation5 Cell Imaging Multi-Mode Reader (BioTek Instruments) was used. All imaging experiments were performed with identical settings. GFP quantifications were performed by using Gen5 3.0 software and via flow vermimetry (Union Biometrica).

### Statistical Analysis

RStudio (version 3.6.3) was used to perform statistical analysis. One-way analysis of variance (ANOVA) was performed to calculate the significance of a treatment when there were three or more groups in the experimental setting. To follow, Dunnett’s test (R package DescTools, version 0.99.34) was performed to calculate statistical significance or *p* values between each group of the statistically significant experimental results. Student’s t test analysis was performed to calculate the *p* values when comparing two groups in an experimental setting. Both Dunnett’s test and Student’s t test results were indicated in graphs as follows: NS not significant, \**p* < 0.05, \*\**p* < 0.01, and ***p < 0.001.

## Acknowledgements

*C. elegans* strains used were provided by David Fay, Cole Haynes, or obtained from the CGC, which is funded by NIH Office of Research Infrastructure Programs (P40 OD010440). We thank Daniel Kirienko for helpful discussion. NVK, a CPRIT scholar in Cancer Research, thanks the Cancer Prevention and Research Institute of Texas (CPRIT) for their generous support, CPRIT grant RR150044. This work was also supported by the National Institutes of Health (NIGMS R35GM129294 to NK.

## Figure 2-source data legend

**Proteomic assays revealed the presence of ESRE-binding factor(s).**

**Source data 1** Raw, unedited **e**lectrophoretic mobility shift assay (EMSA) gel with three tandem ESRE sequences (3XESRE) as a bait.

**Source data 2** Raw, unedited EMSA gel with 3XESRE as a bait, with relevant bands labeled as in **Figure 2B**.

**Source data 3** Raw, unedited EMSA gel with four tandem ESRE sequences (4XESRE) as a bait.

**Source data 4** Raw, unedited EMSA gel with 4XESRE as a bait, with relevant bands labeled as in **Figure 2B**.

---

**Supplementary Figure 1.**
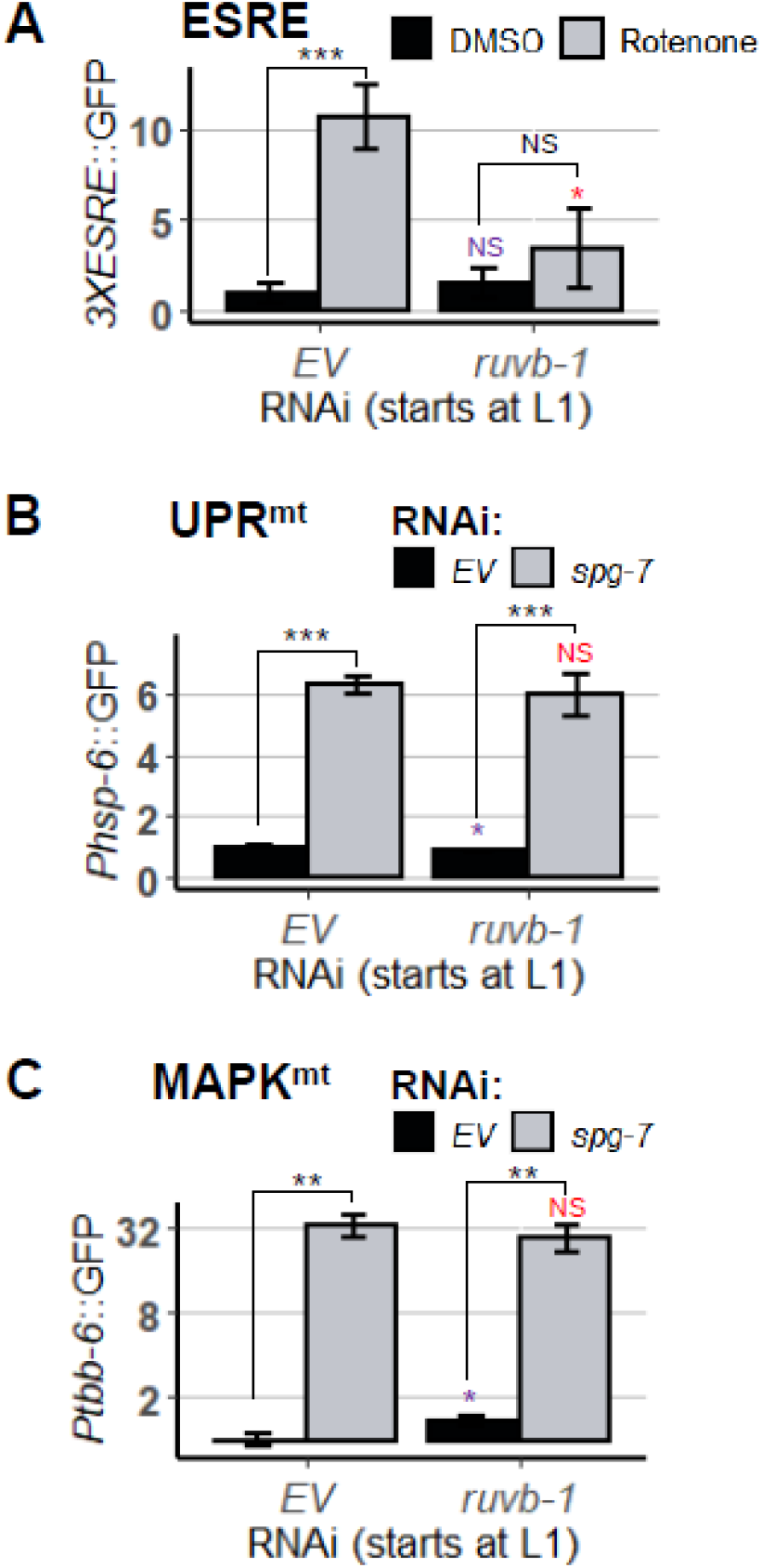
Knockdown of box C/D snoRNP assembly factor RUVB-1 slightly affected the mitochondrial surveillance pathways. Quantification of GFP fluorescence of *C. elegans* carrying **(A)** *3XESRE*::GFP, **(B)** *Phsp-6*::GFP, and **(C)** *Ptbb-6*::GFP reporters that were reared on *E. coli* expressing RNAi targeting empty vector *(EV)* or *ruvb-1/RUVB*. In **(A)**, worms were treated for 8 h with vehicle (DMSO) or 50 μM rotenone. In **(B, C)**, double RNAi was performed with empty vector *(EV)* or *spg-7(RNAi)*. Three biological replicates with ~400 worms/replicate were analyzed. Error bars represent SEM. *p*-values were determined from Student’s *t*-test. GFP values were normalized to EV-DMSO or *EV*. NS not significant, \**p* < 0.05, ** *p* < 0.01, *** *p* < 0.001.

**Supplementary Figure 2.**
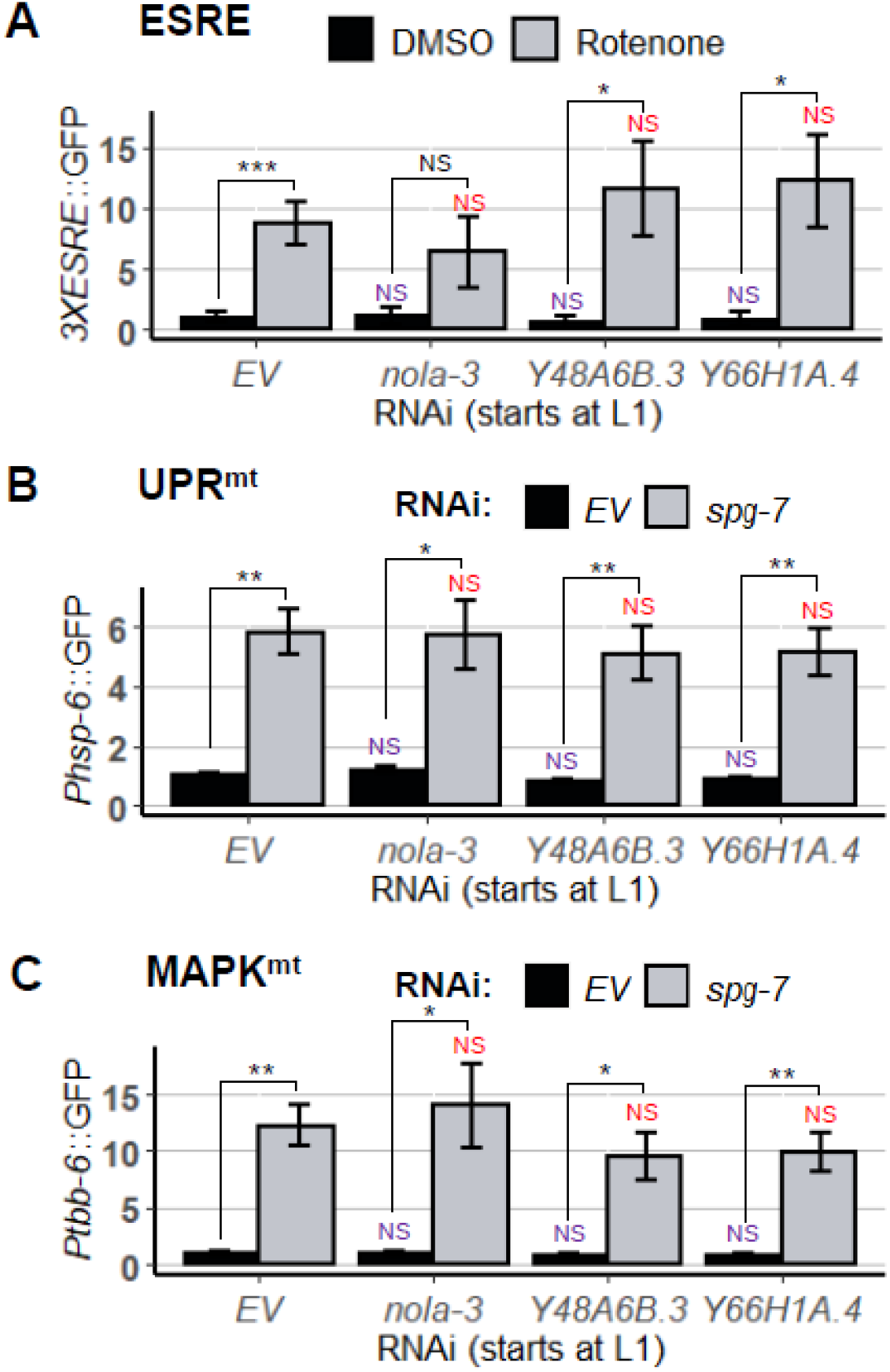
RNAi targeting core members of box H/ACA snoRNPs did not affect mitochondrial surveillance pathways. Quantification of GFP fluorescence of *C. elegans* carrying **(A)** *3XESRE*::GFP, **(B)** *Phsp-6*::GFP, anel **(C)** *Ptbb-6*::GFP reporters that were reared on *E. coli* expressing empty vector *(EV)* or RNAi targeting box H/ACA snoRNP members: *nola-3/Nop10, Y48A68.3/Nhp2*, and *Y66H1A.4/Gar1*. In **(A)**, worms were treated for 8 h with vehicle (DMSO) or 50 μM rotenone. In **(B,C)**, double RNAi was performed with empty vector **(EV)** or *spg-7(RNAi)*. Three biological replicates with ~400 worms/replicate were analyzed. Error bars represent SEM. *p* values were determined from one-way ANOVA, followed by Dunnetl’s test, and Student’s *t*-test. All fold changes were normalized to DMSO-EV or *EV* control. NS not significant, \**p* < 0.05, ** *p* < 0.01, *** *p* < 0.001.

**Supplementary Figure 3.**
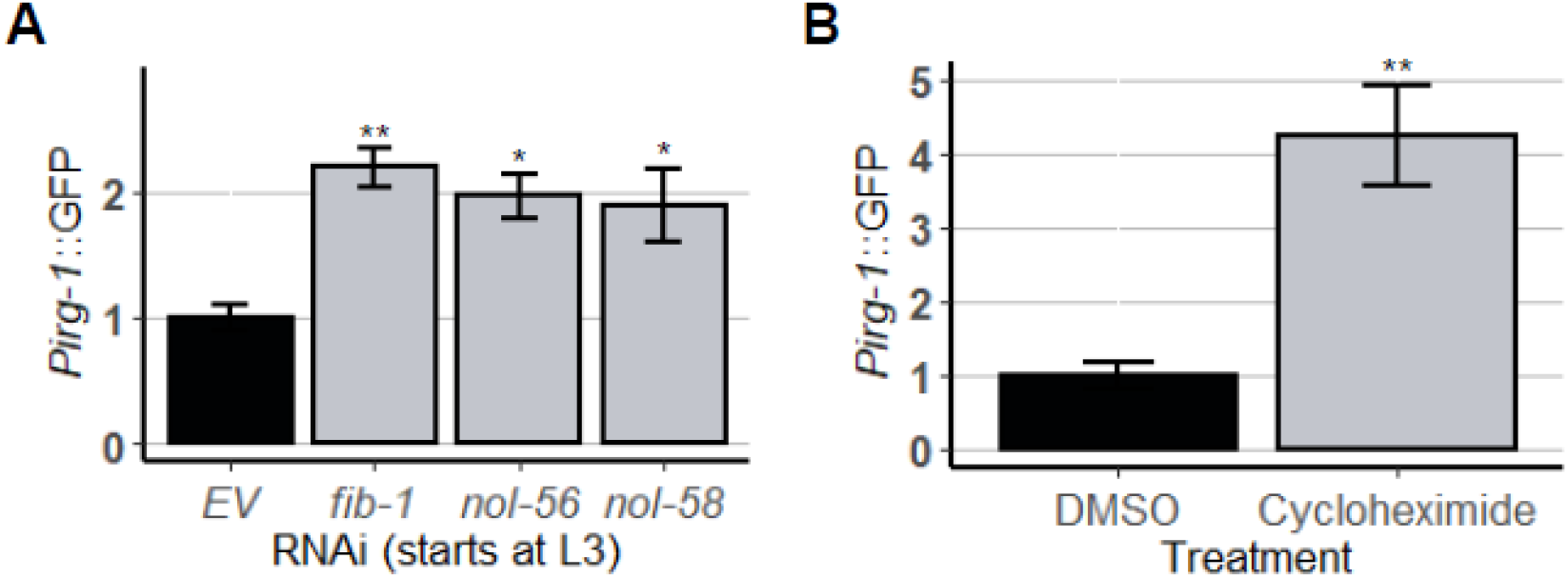
Knockdown of box C/D snoRNPs induced immune response gen *irg-1*. Quantification of GFP fluorescence of *C. elegans* carrying *Pirg-1*::GFP reporter. In **(A)**, worms were reared on *E. coli* expressing empty vector *(EV)*, *fib-1(RNAi), nol-56(RNAi)*, or *nol-58(RNAi)*. In **(B)**. worms were treated for 8 h with vehicle (DMSO) or translation elongation inhibitor cycloheximide. Three biological replicates with ~400 worms/replicate were analyzed. Error bars represent SEM. *p* values were determined from **(A)** one-way ANOVA, followed by Dunnett’s test. or **(B)**, Student’s *t*-test. Fold changes were normalized to **(A)** *EV* control or **(B)** DMSO control. \**p* < 0.05, ** *p* < 0.01.

**Supplementary Figure 4.**
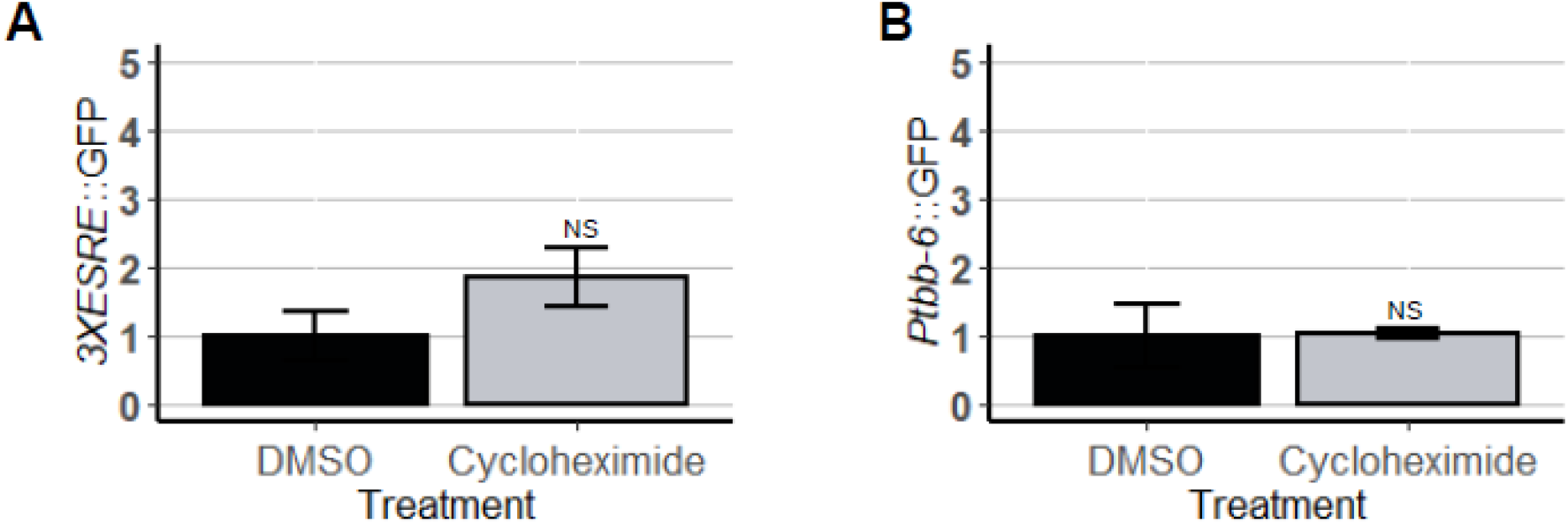
Cycloheximide treatment did not affect mitochondrial surveillance pathways. Quantification of GFP fluorescence of *C. elegans* carrying **(A)** *3XESRE*::GFP or **(B)** *Ptbb-6*::GFP reporters that were treated for 8 h with vehicle (DMSO) or translation elongation inhibitor cycloheximide. Three biological replicates with ~400 worms/replicate were analyzed. Error bars represent SEM. *p* values were determined from Student’s *t*-test. NS not significant.

**Supplementary Figure 5.**
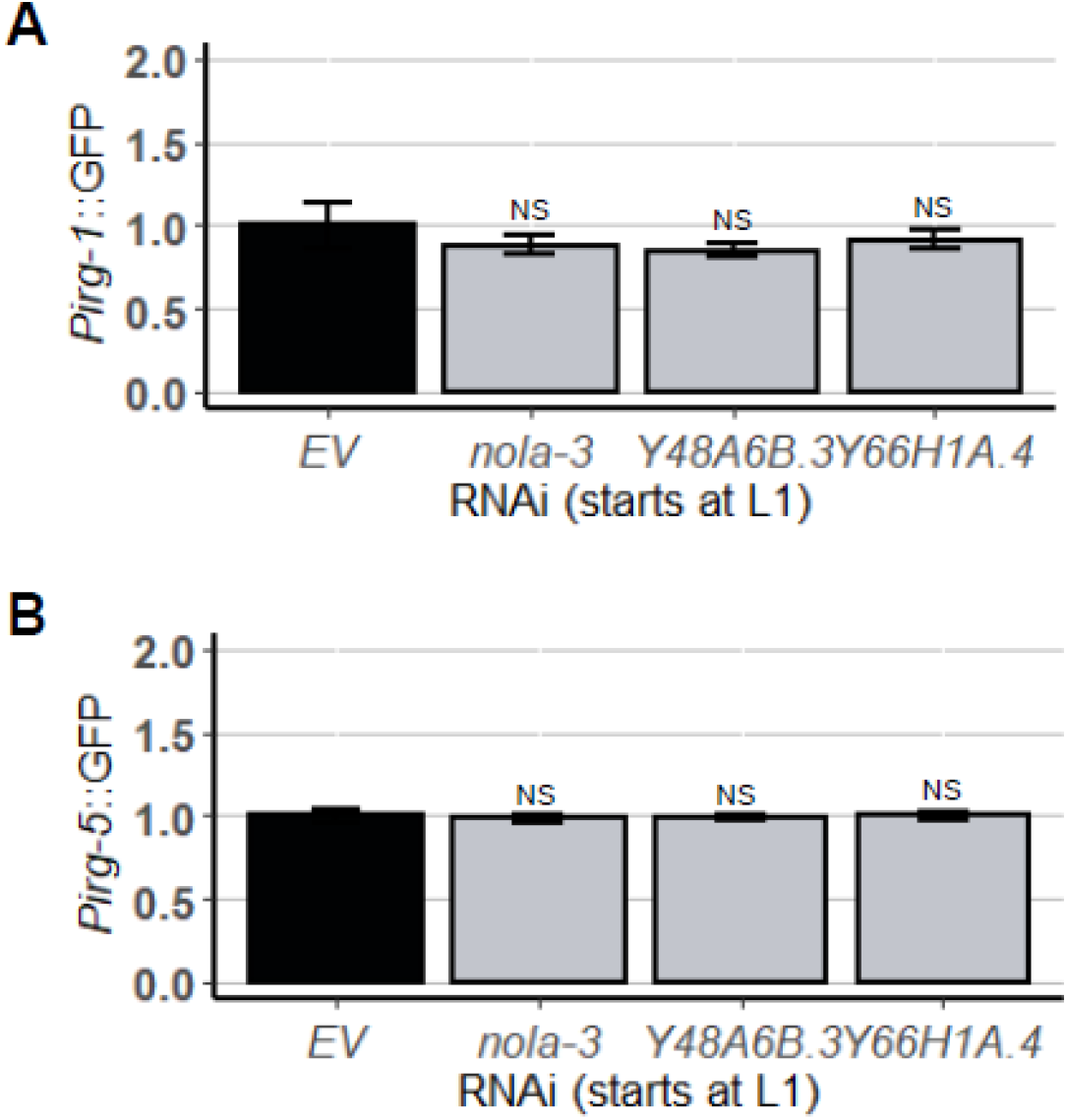
RNAi targeting core members of box H/ACA snoRNPs did not affect innate immune pathways. Quantification of GFP fluorescence of *C. elegans* carrying **(A)** *Pirg-1*::GFP and **(B)** *Pirg-5*::GFP reporters that were reared on *E. coli* expressing empty vector *(EV)* or RNAi targeting box H/ACA snoRNP members: *nola-3/Nop10, Y48A68.31Nhp2*, and *Y66H1A.4/Gar1*. Three biological replicates with ~400 worms/replicate were analyzed. Error bars represent SEM. *p* values were determined from one-way ANOVA, followed by Dunnett’s test. All fold changes were normalized to *EV* control. NS not significant.

## References

1. McEwan Deborah L, Kirienko Natalia V, Ausubel Frederick M. Host Translational Inhibition by Pseudomonas aeruginosa Exotoxin A Triggers an Immune Response in Caenorhabditis elegans. Cell Host & Microbe. 2012;11(4):364–74.

2. Dunbar Tiffany L, Yan Z, Balla Keir M, Smelkinson Margery G, Troemel Emily R. C. elegans Detects Pathogen-Induced Translational Inhibition to Activate Immune Signaling. Cell Host & Microbe. 2012;11(4):375–86.

3. Doye A, Mettouchi A, Bossis G, Clément R, Buisson-Touati C, Flatau G, et al. CNF1 exploits the ubiquitin-proteasome machinery to restrict Rho GTPase activation for bacterial host cell invasion. Cell. 2002;111(4).

4. Garcia-Sanchez JA, Ewbank JJ, Visvikis O. Ubiquitin-related processes and innate immunity in C. elegans. Cell Mol Life Sci. 2021.

5. Jones LM, Chen Y, van Oosten-Hawle P. Redefining proteostasis transcription factors in organismal stress responses, development, metabolism, and health. Biol Chem. 2020;401(9):1005–18.

6. Stradal TEB, Schelhaas M. Actin dynamics in host-pathogen interaction. FEBS Lett. 2018;592(22):3658–69.

7. Alshareef MH, Hartland EL, McCaffrey K. Effectors Targeting the Unfolded Protein Response during Intracellular Bacterial Infection. Microorganisms. 2021;9(4).

8. Choi JA, Song CH. Insights Into the Role of Endoplasmic Reticulum Stress in Infectious Diseases. Front Immunol. 2019;10:3147.

9. Melo Justine A, Ruvkun G. Inactivation of Conserved C. elegans Genes Engages Pathogen- and Xenobiotic-Associated Defenses. Cell. 2012;149(2):452–66.

10. Lemichez E, Barbieri J. General aspects and recent advances on bacterial protein toxins. Cold Spring Harbor perspectives in medicine. 2013;3(2).

11. Khan S, Raj D, Jaiswal K, Lahiri A. Modulation of host mitochondrial dynamics during bacterial infection. Mitochondrion. 2020;53:140–9.

12. Tiku V, Tan MW, Dikic I. Mitochondrial Functions in Infection and Immunity. Trends Cell Biol. 2020;30(4):263–75.

13. Cho DH, Kim JK, Jo EK. Mitophagy and Innate Immunity in Infection. Mol Cells. 2020;43(1):10–22.

14. Bader V, Winklhofer KF. PINK1 and Parkin: team players in stress-induced mitophagy. Biol Chem. 2020;401(6-7):891–9.

15. Youle RJ, Narendra DP. Mechanisms of mitophagy. Nat Rev Mol Cell Biol. 2011;12(1):9–14.

16. Anderson NS, Haynes CM. Folding the Mitochondrial UPR into the Integrated Stress Response. Trends Cell Biol. 2020;30(6):428–39.

17. Fiorese CJ, Haynes CM. Integrating the UPR(mt) into the mitochondrial maintenance network. Crit Rev Biochem Mol Biol. 2017;52(3):304–13.

18. Munkácsy E, Khan MH, Lane RK, Borror MB, Park JH, Bokov AF, et al. DLK-1, SEK-3 and PMK-3 Are Required for the Life Extension Induced by Mitochondrial Bioenergetic Disruption in C. elegans. PLOS Genetics. 2016;12(7):e1006133.

19. Oks O, Lewin S, Goncalves IL, Sapir A. The UPR(mt) Protects Caenorhabditis elegans from Mitochondrial Dysfunction by Upregulating Specific Enzymes of the Mevalonate Pathway. Genetics. 2018;209(2):457–73.

20. Tjahjono E, Kirienko NV. A conserved mitochondrial surveillance pathway is required for defense against Pseudomonas aeruginosa. PLoS Genet. 2017;13(6):e1006876.

21. Tjahjono E, McAnena AP, Kirienko NV. The evolutionarily conserved ESRE stress response network is activated by ROS and mitochondrial damage. BMC Biology. 2020;18(1):74.

22. Kirienko NV, Fay DS. SLR-2 and JMJC-1 regulate an evolutionarily conserved stress-response network. EMBO J. 2010;29(4):727–39.

23. Kwon JY, Hong M, Choi MS, Kang S, Duke K, Kim S, et al. Ethanol-response genes and their regulation analyzed by a microarray and comparative genomic approach in the nematode Caenorhabditis elegans. Genomics. 2004;83(4):600–14.

24. Pignataro L, Varodayan FP, Tannenholz LE, Protiva P, Harrison NL. Brief alcohol exposure alters transcription in astrocytes via the heat shock pathway. Brain Behav. 2013;3(2):114–33.

25. Kirienko NV, Ausubel FM, Ruvkun G. Mitophagy confers resistance to siderophore-mediated killing by Pseudomonas aeruginosa. Proceedings of the National Academy of Sciences. 2015;112(6):1821–6.

26. Kang D, Kirienko DR, Webster P, Fisher AL, Kirienko NV. Pyoverdine, a siderophore from Pseudomonas aeruginosa, translocates into C. elegans, removes iron, and activates a distinct host response. Virulence. 2018;9(1):804–17.

27. Kuzmanov A, Karina EI, Kirienko NV, Fay DS. The Conserved PBAF Nucleosome-Remodeling Complex Mediates the Response to Stress in Caenorhabditis elegans. Molecular and Cellular Biology. 2014;34(6):1121–35.

28. Kirienko NV, McEnerney JD, Fay DS. Coordinated regulation of intestinal functions in C. elegans by LIN-35/Rb and SLR-2. PLoS Genet. 2008;4(4):e1000059.

29. Pukkila-Worley R. Surveillance Immunity: An Emerging Paradigm of Innate Defense Activation in Caenorhabditis elegans. PLoS Pathog. 2016;12(9):e1005795.

30. Anderson S, Cheesman H, Peterson N, Salisbury J, Soukas A, Pukkila-Worley R. The fatty acid oleate is required for innate immune activation and pathogen defense in Caenorhabditis elegans. PLoS pathogens. 2019;15(6).

31. Dasgupta M, Shashikanth M, Gupta A, Sandhu A, De A, Javed S, et al. NHR-49 Transcription Factor Regulates Immunometabolic Response and Survival of Caenorhabditis elegans during Enterococcus faecalis Infection. Infection and immunity. 2020;88(8).

32. Ellis J, Brown D, Brown J. The small nucleolar ribonucleoprotein (snoRNP) database. RNA (New York, NY). 2010;16(4).

33. Ojha S, Malla S, Lyons SM. snoRNPs: Functions in Ribosome Biogenesis. Biomolecules. 2020;10(5):783.

34. Massenet S, Bertrand E, Verheggen C. Assembly and trafficking of box C/D and H/ACA snoRNPs. RNA Biol. 2017;14(6):680–92.

35. Lee J, Harris AN, Holley CL, Mahadevan J, Pyles KD, Lavagnino Z, et al. Rpl13a small nucleolar RNAs regulate systemic glucose metabolism. Journal of Clinical Investigation. 2016;126(12):4616–25.

36. Elliott B, Ho H, Ranganathan S, Vangaveti S, Ilkayeva O, Abou A, H, et al. Modification of messenger RNA by 2’-O-methylation regulates gene expression in vivo. Nature communications. 2019;10(1).

37. Tiku V, Jain C, Raz Y, Nakamura S, Heestand B, Liu W, et al. Small nucleoli are a cellular hallmark of longevity. Nature communications. 2017;8.

38. Tiku V, Kew C, Mehrotra P, Ganesan R, Robinson N, Antebi A. Nucleolar fibrillarin is an evolutionarily conserved regulator of bacterial pathogen resistance. Nature Communications. 2018;9(1):3607.

39. Liang J, Wen J, Huang Z, Chen X, Zhang B, Chu L. Small Nucleolar RNAs: Insight Into Their Function in Cancer. Frontiers in oncology. 2019;9.

40. Mosser D, Kotzbauer P, Sarge K, Morimoto R. In vitro activation of heat shock transcription factor DNA-binding by calcium and biochemical conditions that affect protein conformation. Proceedings of the National Academy of Sciences of the United States of America. 1990;87(10).

41. Saltzman A, Leng M, Bhatt B, Singh P, Chan D, Dobrolecki L, et al. gpGrouper: A Peptide Grouping Algorithm for Gene-Centric Inference and Quantitation of Bottom-Up Proteomics Data. Molecular & cellular proteomics : MCP. 2018;17(11).

42. Lafontaine DL, Tollervey D. Synthesis and assembly of the box C+D small nucleolar RNPs. Mol Cell Biol. 2000;20(8):2650–9.

43. Sheaffer KL, Updike DL, Mango SE. The Target of Rapamycin Pathway Antagonizes pha-4/FoxA to Control Development and Aging. Current Biology. 2008;18(18):1355–64.

44. Ahringer J. Reverse Genetics. WormBook. 2006.

45. Yoneda T, Benedetti C, Urano F, Clark SG, Harding HP, Ron D. Compartment-specific perturbation of protein handling activates genes encoding mitochondrial chaperones. J Cell Sci. 2004;117(Pt 18):4055–66.

46. Nargund AM, Pellegrino MW, Fiorese CJ, Baker BM, Haynes CM. Mitochondrial Import Efficiency of ATFS-1 Regulates Mitochondrial UPR Activation. Science. 2012;337(6094):587–90.

47. Pellegrino MW, Nargund AM, Kirienko NV, Gillis R, Fiorese CJ, Haynes CM. Mitochondrial UPR-regulated innate immunity provides resistance to pathogen infection. Nature. 2014;516(7531):414–7.

48. Erales J, Marchand V, Panthu B, Gillot S, Belin S, Ghayad SE, et al. Evidence for rRNA 2′-O-methylation plasticity: Control of intrinsic translational capabilities of human ribosomes. Proceedings of the National Academy of Sciences. 2017;114(49):12934–9.

49. Penzo M, Montanaro L. Turning Uridines around: Role of rRNA Pseudouridylation in Ribosome Biogenesis and Ribosomal Function. Biomolecules. 2018;8(2).

50. Kavčič B, Tkačik G, Bollenbach T. Mechanisms of drug interactions between translation-inhibiting antibiotics. Nature communications. 2020;11(1).

51. Rodnina M. The ribosome in action: Tuning of translational efficiency and protein folding. Protein science : a publication of the Protein Society. 2016;25(8).

52. Bolz DD, Tenor JL, Aballay A. A Conserved PMK-1/p38 MAPK Is Required in Caenorhabditis elegans Tissue-specific Immune Response to *Yersinia pestis* Infection. Journal of Biological Chemistry. 2010;285(14):10832–40.

53. Pukkila-Worley R, Feinbaum R, Kirienko NV, Larkins-Ford J, Conery AL, Ausubel FM. Stimulation of Host Immune Defenses by a Small Molecule Protects C. elegans from Bacterial Infection. PLoS Genetics. 2012;8(6):e1002733.

54. Shivers RP, Pagano DJ, Kooistra T, Richardson CE, Reddy KC, Whitney JK, et al. Phosphorylation of the Conserved Transcription Factor ATF-7 by PMK-1 p38 MAPK Regulates Innate Immunity in Caenorhabditis elegans. PLoS Genetics. 2010;6(4):e1000892.

55. Peterson N, Cheesman H, Liu P, Anderson S, Foster K, Chhaya R, et al. The nuclear hormone receptor NHR-86 controls anti-pathogen responses in C. elegans. PLoS genetics. 2019;15(1).

56. Kirienko Natalia V, Kirienko Daniel R, Larkins-Ford J, Wählby C, Ruvkun G, Ausubel Frederick M. Pseudomonas aeruginosa Disrupts Caenorhabditis elegans Iron Homeostasis, Causing a Hypoxic Response and Death. Cell Host & Microbe. 2013;13(4):406–16.

57. Kang D, Kirienko NV. An In Vitro Cell Culture Model for Pyoverdine-Mediated Virulence. Pathogens. 2020;10(1).

58. Kang D, Revtovich AV, Chen Q, Shah KN, Cannon CL, Kirienko NV. Pyoverdine-Dependent Virulence of Pseudomonas aeruginosa Isolates From Cystic Fibrosis Patients. Front Microbiol. 2019;10:2048.

59. Feinbaum RL, Urbach JM, Liberati NT, Djonovic S, Adonizio A, Carvunis AR, et al. Genome-wide identification of Pseudomonas aeruginosa virulence-related genes using a Caenorhabditis elegans infection model. PLoS Pathog. 2012;8(7):e1002813.

60. Lee DG, Urbach JM, Wu G, Liberati NT, Feinbaum RL, Miyata S, et al. Genomic analysis reveals that Pseudomonas aeruginosa virulence is combinatorial. Genome Biol. 2006;7(10):R90.

61. Kim DH, Feinbaum R, Alloing G, Emerson FE, Garsin DA, Inoue H, et al. A conserved p38 MAP kinase pathway in Caenorhabditis elegans innate immunity. Science. 2002;297(5581):623–6.

62. Kim DH, Liberati NT, Mizuno T, Inoue H, Hisamoto N, Matsumoto K, et al. Integration of Caenorhabditis elegans MAPK pathways mediating immunity and stress resistance by MEK-1 MAPK kinase and VHP-1 MAPK phosphatase. Proc Natl Acad Sci U S A. 2004;101(30):10990–4.

63. Troemel ER, Chu SW, Reinke V, Lee SS, Ausubel FM, Kim DH. p38 MAPK regulates expression of immune response genes and contributes to longevity in C. elegans. PLoS Genet. 2006;2(11):e183.

64. Hummell NA, Revtovich AV, Kirienko NV. Novel Immune Modulators Enhance Caenorhabditis elegans Resistance to Multiple Pathogens. mSphere. 2021;6(1).

65. Vitali P, Basyuk E, Le Meur E, Bertrand E, Muscatelli F, Cavaille J, et al. ADAR2-mediated editing of RNA substrates in the nucleolus is inhibited by C/D small nucleolar RNAs. J Cell Biol. 2005;169(5):745–53.

66. Vitali P, Kiss T. Cooperative 2’-O-methylation of the wobble cytidine of human elongator tRNA(Met)(CAT) by a nucleolar and a Cajal body-specific box C/D RNP. Genes Dev. 2019;33(13-14):741–6.

67. Aw JG, Shen Y, Wilm A, Sun M, Lim XN, Boon KL, et al. In Vivo Mapping of Eukaryotic RNA Interactomes Reveals Principles of Higher-Order Organization and Regulation. Mol Cell. 2016;62(4):603–17.

68. Sharma E, Sterne-Weiler T, O’Hanlon D, Blencowe BJ. Global Mapping of Human RNA-RNA Interactions. Mol Cell. 2016;62(4):618–26.

69. Holley CL, Li MW, Scruggs BS, Matkovich SJ, Ory DS, Schaffer JE. Cytosolic accumulation of small nucleolar RNAs (snoRNAs) is dynamically regulated by NADPH oxidase. J Biol Chem. 2015;290(18):11741–8.

70. Chen MS, Goswami PC, Laszlo A. Differential accumulation of U14 snoRNA and hsc70 mRNA in Chinese hamster cells after exposure to various stress conditions. Cell Stress Chaperones. 2002;7(1):65–72.

71. Mleczko AM, Machtel P, Walkowiak M, Wasilewska A, Pietras PJ, Bakowska-Zywicka K. Levels of sdRNAs in cytoplasm and their association with ribosomes are dependent upon stress conditions but independent from snoRNA expression. Sci Rep. 2019;9(1):18397.

72. Sloan KE, Warda AS, Sharma S, Entian KD, Lafontaine DLJ, Bohnsack MT. Tuning the ribosome: The influence of rRNA modification on eukaryotic ribosome biogenesis and function. RNA Biol. 2017;14(9):1138–52.

73. Stiernagle T. Maintenance of C. elegans. WormBook. 2006:1–11.

74. Rauthan M, Ranji P, Aguilera Pradenas N, Pitot C, Pilon M. The mitochondrial unfolded protein response activator ATFS-1 protects cells from inhibition of the mevalonate pathway. Proceedings of the National Academy of Sciences. 2013;110(15):5981–6.

75. Kamath RS, Fraser AG, Dong Y, Poulin G, Durbin R, Gotta M, et al. Systematic functional analysis of the Caenorhabditis elegans genome using RNAi. Nature. 2003;421(6920):231–7.

76. Rual J-F, Ceron J, Koreth J, Hao T, Nicot A-S, Hirozane-Kishikawa T, et al. Toward improving Caenorhabditis elegans phenome mapping with an ORFeome-based RNAi library. Genome research. 2004;14(10b):2162–8.

77. Mathee K. Forensic investigation into the origin of Pseudomonas aeruginosa PA14 – old but not lost. J Med Microbiol. 2018;67(8):1019–21.

78. Anderson QL, Revtovich AV, Kirienko NV. A High-throughput, High-content, Liquid-based C. elegans Pathosystem. Journal of visualized experiments : JoVE. 2018(137).

79. Kirienko NV, Cezairliyan BO, Ausubel FM, Powell JR. Pseudomonas aeruginosa PA14 pathogenesis in Caenorhabditis elegans. Methods Mol Biol. 2014;1149:653–69.

